# Development of a novel VHH intrabody targeting the N17 region of huntingtin exon 1 protein that prevents inclusion body formation

**DOI:** 10.64898/2026.04.09.716913

**Authors:** Florence D.M. Wavreil, Wouter Pos, Menno Spits, Alicia Sanz Sanz, Kes Rietveld, Roos van Dam, Martino Böhne, Sander van Deventer, Sabine Schipper-Krom, Eric A.J. Reits

**Affiliations:** Amsterdam UMC, Department of Medical Biology, Amsterdam, The Netherlands; VectorY Therapeutics B.V., Amsterdam, The Netherlands

**Keywords:** Huntington’s disease, Huntingtin exon 1, VHH, Intrabody, Inclusion bodies, Aggregation

## Abstract

Huntington’s disease (HD) is a progressive neurodegenerative disease caused by a mutation in the exon 1 of the huntingtin (*HTT*) gene, which leads to an extended polyglutamine (polyQ) tract in the mutant protein. As a result, mutant huntingtin (mHTT) exon 1 fragments aggregate in cells, which disrupts proper neuronal function and eventually induces cell death. The selective reduction of these toxic mHTT fragments without disturbing the wild-type full-length HTT function would be a potential therapeutic strategy to treat and prevent HD. Intracellular antibodies (intrabodies) have emerged as an attractive strategy to specifically target disease-related proteins, with VHH intrabodies being of high interest as they are much smaller than single-chain variable fragments (scFv). Here, we describe the identification and development of VHH 1 as a lead candidate intrabody targeting the first 17 amino acids of the mHTT protein, using a humanized VHH page-display library to screen against mHTT(Q46) exon 1 to identify potential binders. Next, we further optimized VHH 1 into VHH 1a to improve cytoplasmic solubility. Using immortalized mouse striatal cells that express inducible untagged mHTT exon 1 fragments, we investigated the effects of the intrabody on soluble and insoluble mHTT species via microscopy and biochemical assays. We showed that the VHH 1a intrabody reduces the levels of insoluble mHTT species, thereby effectively interrupting the aggregation process. This study highlights the potential for VHH intrabodies to specifically target mHTT fragments, enabling therapeutic strategies to delay and prevent HD pathology.

**Highlights:**

- Three binders were down-selected from a phage-display library to bind HTT N17
- VHH 1a intrabody is the most efficient at reducing mutant HTT exon 1 aggregation
- VHH 1a acts on soluble HTT exon 1 oligomers to block the transition to inclusion body

## Introduction

Huntington’s disease (HD) is an autosomal-dominant, progressive neurodegenerative disorder characterized by motor disorders such as chorea as well as cognitive decline, behavioral changes, and psychiatric manifestations (Tabrizi et al., 2020). The disease typically manifests around 40 years of age, and as it advances, affected individuals experience a marked reduction in life quality as well as an increased dependence on nursing care and a shortened lifespan (McColgan & Tabrizi, 2018). No disease-modifying treatments exist for HD, underscoring the need to develop new therapeutic strategies to prevent or reverse HD pathology.

The disease is caused by a mutation in the exon 1 of the huntingtin (*HTT*) gene, which results in a CAG trinucleotide repeat expansion and an extended polyglutamine (polyQ) tract at the N-terminus of the mutant HTT (mHTT) protein. The length of the CAG repeat is a key determinant of disease penetrance and age of onset with longer repeats correlating with earlier onset and more severe pathology (Andrew et al., 1993; Consortium, 2019; Duyao et al., 1993; Swami et al., 2009).

Wild-type HTT plays essential roles in neuronal function including axonal transport, transcriptional regulation, and mitochondrial homeostasis (Liu & Zeitlin, 2017; Schulte & Littleton, 2011). Hence, disrupting the protein activity can be detrimental, eventually leading to cell death. Although HTT is ubiquitously expressed, mHTT primarily affects striatal medium spiny neurons and, to a lesser extent, cortical neurons (Han et al., 2010). The susceptibility of the neurons is thought to arise from somatic hypermutation of the *HTT* gene, which mainly occurs in striatal medium spiny neurons and causes the CAG tract to expand up to hundreds of CAG repeats (Handsaker et al., 2025). The increase in CAG expansions subsequently promotes aberrant splicing of mHTT, generating the toxic N-terminal exon 1 (HTT1a) fragments (Papadopoulou et al., 2025; Sathasivam et al., 2013). Exon 1 fragments are believed to be more toxic than the full-length protein and to be sufficient to initiate HD phenotypes (Ross & Tabrizi, 2011; van der Bent et al., 2022). Indeed, exon 1 fragments are prone to misfolding and aggregation as they can undergo a transition from soluble species (monomer and small oligomers) to insoluble species such as aggregates and large inclusion bodies (Legleiter et al., 2010; Leitman et al., 2013; Nucifora et al., 2012).

One promising therapeutic strategy for HD involves selectively reducing the levels of toxic mHTT exon 1 species while preserving the full-length HTT function. Antibody-based approaches have emerged as an attractive strategy to target pathological proteins (Denis et al., 2019). Specifically, intracellular antibodies (intrabodies) are engineered antibody fragments that act within cells, where they can bind, sequester, neutralize, or promote the degradation of target proteins. Intrabodies are most commonly developed in one of three forms: single-chain variable fragments (scFv) that contain the heavy (VH) and light (VL) variable regions of an antibody fused by a linker, single-domain antibodies consisting of only the VH or VL domain, and VHH intrabodies which consist of the variable domain of a heavy chain-only camelid antibody. Compared to scFv, VHHs are expected to be much more stable in the cytosolic environment yet equally efficient at targeting pathogenic proteins (Soetens et al., 2020). Furthermore, their single-domain structure reduces the risk of self-aggregation, and their smaller size (∼15 kDa) relative to scFv (∼35 kDa) is an advantage for the delivery of the intrabody as gene therapy (Asaadi et al., 2021). Several scFv intrabodies targeting different regions of HTT proteins have previously been developed with varying degrees of efficacy (Denis et al., 2019; Jurcau et al., 2024). For example, earlier work demonstrated that a scFv intrabody (C4) directed against the N-terminal region of mHTT could reduce aggregation and toxicity in cellular, mouse, and fly models (Butler & Messer, 2011; Lecerf et al., 2001; Murphy & Messer, 2004; Snyder-Keller et al., 2010), underscoring the therapeutic potential of intrabodies in HD.

In this study, we report the development of a novel VHH intrabody targeting the N-terminus of mHTT exon 1. Within this work, a humanized VHH phage-display library screened against mHTT(Q46) exon 1 fragment identified the VHH 1 intrabody, which binds the N-terminal 17 amino acids of HTT. Via two point mutations, we further optimized the VHH 1 intrabody to improve solubility in the cytosolic environment. We then evaluated the effects of the VHHs on soluble and insoluble untagged mHTT exon 1 protein fragments in mouse striatal cell lines via a combination of microscopy and biochemical assays.

## Materials and Methods

### VHH single domain antibody discovery and down-selection

Humanized single-domain antibodies against HTT exon 1 were identified using a humanized VHH phage display library (∼5×10⁸ unique sequences) (Alexander & Leong, 2024). The library was screened against multiple HTT exon 1 antigen formats to ensure broad epitope coverage. Antigens included synthetic peptides (BioSynth) corresponding to N17 domain, polyglutamine (PolyQ), proline-rich region (PRR), and C-terminal regions, and recombinant HIS-MBP-HTT exon 1 proteins with 25 (Q25) or 46 (Q46) glutamines produced by Trenzyme. Four independent panning arms enriched for linear and conformational binders, incorporating competitive deselection to eliminate non-specific and MBP-directed interactions. Four rounds of panning were conducted under increasing stringency, followed by Sanger sequencing for clone identification. Primary screening used AlphaLISA (Revvity) (Eglen et al., 2008) to detect binding to biotinylated N17, PolyQ, and PRR peptides or HIS-MBP fusion proteins. Clones with a signal-to-noise ratio above 25 were classified as binders. Positive clones were then characterized by surface plasmon resonance (SPR) (Hojjat Jodaylami et al., 2025) to assess affinity and specificity for MBP-tagged HTT exon 1 Q25 and Q46 variants. C-terminally V5-tagged VHHs were captured on α-V5 sensor chips, and binding kinetics were analyzed using a 1:1 Langmuir model to determine KD values.

### Plasmid construction

The transgenes for selected intrabodies, both VHH and scFv formats which included positive and negative controls, were cloned by Azenta into an already available plasmid containing the CBh promoter and SV40 polyA sequences. Plasmids were purified and delivered according to the manufacturer’s protocol.

### Peptide synthesis

Synthetic peptides representing N17, PolyQ, PolyP, PRR, and C-terminal regions of HTT exon 1 were produced by BioSynth B.V. (Lelystad, The Netherlands) using standard Fmoc solid-phase peptide synthesis. Each peptide carried a C-terminal biotin tag for immobilization, with a GGGS linker included when necessary to reduce steric hindrance. To map the epitope and assess antibody specificity and sensitivity to modification, alanine-substituted and post-translationally modified N17 variants were synthesized, including acetylation (K6, K9, K15), phosphorylation (T3, S13, S16), and M1 truncation, each verified by LC–MS.

### Recombinant protein expression and purification

Humanized VHHs for ELISA measurements were expressed in Expi293F cells for proper folding and post-translational modification. The VHH-His format included a C-terminal His tag, and the VHH-IgG1 format fused the VHH to a human IgG1 Fc for enhanced expression and bivalency. Cells were transiently transfected using polyethyleneimine and cultured in serum-free medium for 5–7 days. hVHH-His proteins were purified by Ni-NTA affinity chromatography and eluted with imidazole. hVHH-IgG1 fusions were purified by Protein A chromatography and buffer-exchanged into PBS. Protein aggregates were removed by size exclusion chromatography in PBS. Purity was confirmed by SDS-PAGE, and concentration was determined by UV absorbance at 280 nm. A non-targeting control VHH (Ashour et al., 2015) was included for assay validation.

Recombinant MBP-tagged HTT exon 1 proteins with 25 (Q25) or 46 (Q46) glutamines were produced by Trenzyme GmbH (Konstanz, Germany) in HEK293 cells to ensure proper folding and processing. Proteins were purified by IMAC and desalted into PBS. Quality and purity (>99%) were confirmed by SDS-PAGE and Coomassie staining, and identity was verified by western blot with rabbit α-HTT EPR5526 (Abcam, ab109115). No size-exclusion step was applied due to aggregation and yield constraints.

### Peptide and recombinant protein binding ELISAs

Binding of single-domain VHHs (VHH-His) and IgG1-reformatted VHHs (VHH-IgG1) was assessed by peptide-based ELISA to confirm specificity for HTT exon 1 regions. Biotinylated peptides were immobilized on streptavidin-coated 96-well plates. Plates were blocked with phosphate-buffered saline containing 0.1% Tween-20 (PBST) and 3% bovine serum albumin (BSA). VHH-His and VHH-IgG1 antibodies were diluted in the blocking buffer and incubated for one hour at room temperature. Detection used an unconjugated α -HIS primary antibody followed by HRP-labeled α-mouse IgG, which produced optimal signal and specificity for VHH-His samples. After washing with PBST, bound HRP was detected using 3,3′,5,5′-tetramethylbenzidine (TMB) substrate and stopped with 1 M H₂SO₄. Absorbance was read at 450 nm. All assays were performed in triplicate and included an irrelevant scFv as a negative control. Signal-to-noise ratios were calculated to assess assay performance.

Binding of VHH-His and VHH-hIgG1 formats to recombinant MBP-HTT exon 1 proteins with 25 (Q25) or 46 (Q46) glutamines was measured by ELISA. High-binding plates were coated with purified MBP-HTT exon 1 Q25 or Q46 proteins and blocked with 3% BSA in PBST. Bound VHHs were detected using α-HIS-HRP for His-tagged constructs or α-human IgG-HRP for Fc-fused constructs. Absorbance was measured at 450 nm after the addition of the TMB substrate and the stop solution. All assays were run in duplicate and validated using independent antigen batches.

### Cell culture and transfection

Human U2OS cells were cultured at 37°C with 5% CO_2_ in complete Dulbecco’s Modified Eagle Medium (DMEM; Gibco) supplemented with 10% fetal bovine serum (FBS), 1% penicillin/streptomycin, and 2µg/mL puromycin. Mouse striatal STHdh cells were cultured at 32°C with 5% CO_2_ in complete DMEM supplemented with 10% FBS, 1% penicillin/streptomycin, and 0.2 mM L-glutamine. Stable STHdh^Q7/Q7^ cells expressing N-terminal HTT exon 1 fragment with various glutamine repeats under a doxycycline inducible promoter were previously generated by lentiviral transduction with pINDUCER mHTT(Q25/Q97)-IRES-GFP-Q16 (Geijtenbeek et al., 2022). U2OS cells expressing doxycycline-inducible mHTT(Q97)-IRES-GFPQ16 were generated as previously described for the STHdh cells (Geijtenbeek et al., 2022).

U2OS cells were seeded in a poly-D-lysine coated 96-well plate and treated with 1μg/mL of doxycycline (Sigma-Aldrich) to induce mHTT expression. The next day, cells were transfected with FLAG-tagged intrabody constructs (50 ng DNA/8,000 cells) using Lipofectamine 3000 reagent according to the manufacturer’s instructions. Cells were fixed 72 hours after seeding. STHdh cells were transfected with Flag-tagged intrabody constructs (250 ng DNA/100,000 cells) via electroporation using the Neon^TM^ Electroporation System (Invitrogen) with two 20 ms pulses of 1300 V according to the manufacturer’s instructions. Depending on the experiment, either 4×10^5^ or 4×10^6^ cells were electroporated with the 10 µL or 100 µL Neon^TM^ kit, respectively, before being seeded into a 6-well plate or a 10 cm dish. HTT exon 1 fragment expression was induced with 1μg/mL of doxycycline either directly upon electroporation or 48 hours prior. Cells were harvested or fixed 72 hours post-electroporation.

### Microscopy aggregate quantification

Cells were plated in a 96-well plate and fixed for 15 minutes with 4% paraformaldehyde (U2OS cells) or 4% formaldehyde (STHdh cells) 72 hours after transfection. U2OS cells were permeabilized with 0.2% Triton-X-100 and blocked with 3% BSA before staining with rabbit α-HTT EPR5526 (1:1000; Abcam, ab109115) and mouse α-FLAG (1:800; Sigma, F3165) primary antibodies followed by α-rabbit Alexa Fluor 647 (1:200; Invitrogen, A21245) and α-mouse Alexa Fluor 568 (1:200; Invitrogen, A11004) secondary antibodies. Nuclei were stained with DAPI. High-content imaging was conducted on an ImageXpress Micro Confocal system (20x objective). Nine fields per well were captured. The number of aggregates per FLAG-positive cells was quantified using MetaXpress software with automated segmentation of nuclei, cytoplasm, and aggregates.

STHdh cells were permeabilized for 1 hour (2% FBS, 1% BSA, 0.4% Triton-X-100 in PBS), then stained with mouse α-FLAG (1:800; Sigma, F3165) primary antibody overnight at 4°C and α-mouse Cy3 (1:200; Jackson ImmunoResearch, 115-165-166) secondary antibody for 1 hour at room temperature. Nuclei were stained with 5 μg/ml Hoechst dye 33324 (Invitrogen, H3570) for 1 hour at room temperature. Images were acquired with an automated widefield microscope (20x objective; ImageXpress Pico). The percentage of FLAG-positive cells with aggregates was analyzed based on a modified MATLAB algorithm previously developed (Geijtenbeek et al., 2022). Briefly, the number of cells is determined based on nuclear segmentation of the Hoechst staining, the number of aggregates is quantified based on the GFP-Q16 fluorescent reporter, and the transfection efficiency is determined based on the number of cells showing a FLAG signal intensity that is 4 times the median absolute deviation (MAD) above that of the negative control.

### Immunofluorescence staining and confocal imaging

Cells were plated on glass coverslips in a 12-well plate and fixed for 15 minutes with 4% formaldehyde 72 hours after electroporation. After 1 hour permeabilization (2% FBS, 1% BSA, 0.4% Triton-X-100 in PBS), cells were stained with rabbit α-HTT EPR5526 (1:1000; Abcam, ab109115) and mouse α-FLAG (1:800; Sigma, F3165) primary antibodies overnight at 4°C. Secondary antibody staining was done at room temperature for 1 hour with α-rabbit Alexa Fluor 633 (1:200; Invitrogen, A21071) and α-mouse Cy3 (1:200; Jackson ImmunoResearch, 115-165-166) antibodies. Coverslips were mounted on glass slides using a drop of Vectashield mounting medium containing DAPI (Vector Laboratories, H-1200). Samples were imaged on a Leica Stellaris 5 confocal microscope using a 63x/1.40 oil immersion objective.

### SDS-PAGE and western blot

Cell pellets were lysed in 1% Triton-X-100 lysis buffer (50 mM Tris/HCl pH 7.4, 150 mM NaCl, 1% Triton-X-100, supplemented with complete mini protease inhibitor cocktail (Roche)) and incubated for 30 minutes on ice. Samples were centrifuged (15 minutes, 4°C at 14000 rpm) to isolate the supernatant, whereas the pellets were saved for filter trap assay. Protein concentration was determined by Bradford assay (Serva). Sample volumes representing equal protein levels were boiled at 99°C for 5 minutes at 750 rpm in 6x sample loading buffer (350 mM Tris/HCl pH 6.8, 10% SDS, 30% glycerol, 6% β-mercaptoethanol, bromophenol blue). Proteins were separated at 120 V on a 12.5% SDS-PAGE gel for HTT exon 1 or on a 4-15% gradient gel (Bio-Rad, 456-1084) for full-length HTT in running buffer (25 mM Tris base, 192 mM glycine, 0.1% SDS) before being transferred for 30 minutes in transfer buffer (20% ethanol, 1x Trans-Blot Turbo buffer (Bio-Rad, 10026938)) to a nitrocellulose membrane (Bio-Rad) using the Trans-Blot Turbo Transfer System (Bio-Rad). Membranes were blocked in 5% milk for 1 hour at room temperature, then incubated overnight at 4°C with primary antibodies and 1 hour at room temperature with secondary antibodies. Primary antibodies used included rabbit α-HTT EPR5526 (1:3000; Abcam, ab109115), mouse α-βactin (1:2000; Santa Cruz Biotechnology, SC-47778), mouse α-vinculin (1:1000; Merck, V9131), and mouse α-FLAG (1:800; Sigma, F3165). Secondary antibodies included IRDye 680RD α-mouse (LI-COR Biosciences, 926-68072) at 1:10000 and IRDye 800CW α-rabbit (LI-COR Biosciences, 926-32213) at 1:10000. Membranes were imaged and analyzed using the Odyssey imaging system (LI-COR Biosciences). The HTT levels were normalized to either actin or vinculin before being normalized to the VHH control.

### Filter trap assay

Pellets obtained after cell lysis and centrifugation were resuspended and incubated for 1 hour at 37°C in endonuclease buffer (1 mM MgCl_2_, 50 mM Tris/HCl pH 8.0, with 0.02 U/μL DENARASE® (c-LEcta) added fresh). The endonucleases activity was stopped using 2x termination buffer (40 mM EDTA, 4% SDS, 100 mM DTT fresh). Samples containing 30μg of protein were prepared in 2% SDS buffer (2% SDS, 150 mM NaCl, 10 mM Tris/HCl pH8.0). Using the Bio-Dot microfiltration system (Bio-Rad, Hercules), samples were loaded in duplicate and filtered through a 0.2 μm cellulose acetate membrane (Schleicher & Schuell) pre-equilibrated in 2% SDS buffer. The membrane was then washed twice with 0.1% SDS buffer (0.1% SDS, 150 mM NaCl, 10 mM Tris pH 8.0). The membrane was blocked in 5% milk, incubated with antibodies, and analyzed as described for the western blot membranes. Antibodies used included rabbit α-HTT EPR5526 (1:3000; Abcam, ab109115) and IRDye 800CW α-rabbit (1:10000; LI-COR Biosciences, 926-32213). For the analysis, the duplicates were averaged before being normalized to the VHH control.

### Agarose Gel Electrophoresis for Resolving Aggregates (AGERA)

Cell pellets were lysed for 30 minutes on ice in 1% Triton-X-100 lysis buffer (50 mM Tris/HCl pH 7.4, 150 mM NaCl, 1% Triton-X-100, supplemented with complete mini protease inhibitor cocktail (Roche)). The lysate was then sonicated (40% amplitude, 40” ON 20” OFF for 13min20) using Qsonica Q800R2 sonicator. Protein concentration was determined by Bradford assay (Serva), and samples containing 70 μg of protein were prepared in 4x Laemmli buffer (Bio-Rad, 161-0747). Samples were run on a 2% agarose gel (2 g agarose in 100 mL 1x TAE and 0.1% SDS) kept on ice at 100V for about 1h45 in running buffer (1X TAE, 0.1% SDS). Proteins were transferred through capillary blotting onto a pre-activated PVDF membrane (Immobilon FL, IPFL00010) for 24 hours using methanol-containing transfer buffer (25 mM Tris base, 192 mM glycine, 0.1% SDS, 15% methanol). The membrane was then blocked and treated as described for the western blot membrane. Antibodies used included sheep α-HTT S830 (1:1000, kindly provided by Prof. G. Bates, University College, London, UK) and IRDye 680RD α-goat (1:10000; LI-COR Biosciences, 926-68074). After drawing a rectangular region of interest around each smear, the total signal intensity of the smears and a density profile plot for each sample were analyzed with ImageJ.

### Blue Native Polyacrylamide Gel Electrophoresis (BN-PAGE)

Cell pellets were lysed for 30 minutes on ice in 1% Triton-X-100 lysis buffer (20 mM Bis-Tris pH 7.0, 20 mM NaCl, 10% glycerol, 0.1% Triton-X-100, supplemented with complete mini protease inhibitor cocktail (Roche)). Samples were centrifuged briefly (5 minutes, 4°C at 14000 rpm), and the supernatant was isolated. Protein concentration was determined with a Bradford assay (Serva), and samples containing 35μg of protein were prepared in 4x loading buffer (20 mM Bis-Tris pH 7.0, 20 mM NaCl, 40% glycerol, 0.5% Coomassie Blue G250). Samples were run at 150 V on a 3-15% Bis-Tris Native PAGE gel (Invitrogen, BN2012BX10) using Bis-Tris anode buffer (50 mM Bis-Tris pH 7.0) and Tricine cathode buffer (50 mM Tricine, 15 mM Bis-Tris pH 7.0). For the first 20 minutes of running, the cathode buffer was supplemented with 0.01% Coomassie Brilliant Blue G250 before being replaced by fresh cathode buffer. Proteins were transferred onto a pre-activated PVDF membrane (Immobilon FL, IPFL00010) for 30 minutes in transfer buffer (20% ethanol, 1x Trans-Blot Turbo buffer (Bio-Rad, 10026938)) using the Trans-Blot Turbo Transfer System (Bio-Rad). After the transfer, the membrane was washed twice with methanol to remove the excess Coomassie Brilliant blue G250 and then blocked in 5% milk and treated as described for the western blot membrane. Antibodies used included sheep α-HTT S829 (1:1000, kindly provided by Prof. G. Bates, University College, London, UK) and IRDye 680RD α-goat (1:10000; LI-COR Biosciences, 926-68074).

### Cell viability assay

Cells were plated in triplicate in a 96-well plate at a density of 10000 cells/well. After 72 hours post-electroporation, the culture medium was removed, and cells were incubated with 3-(4,5-dimethylthiazol-2-yl)-2,5-diphenyltetrazolium-bromide (MTT; Promega, G402A) diluted 1:10 in complete DMEM for 4 hours at 32°C. Next, 100 μL of solubilization solution (Promega, G401A) was added to each well, and cells were incubated for 1 hour at 32°C. Absorbance was read on a CLARIOstar plate reader (BMG LABTECH) at 570 nm and at 690nm as reference wavelength. For the analysis, the absorbance values at 690 nm were subtracted from the 570 nm absorbance values. The triplicate values were averaged and normalized to the VHH control.

### Data quantification and statistics

Quantitative data were analyzed using GraphPad Prism 10 software. Results are presented as mean ± SD. Outliers were identified using the ROUT test (Q = 10%) and removed. Statistical differences between groups were identified by a one-way ANOVA with Tukey test for multiple comparisons. An alpha level of 0.05 was used to define statistical significance (ns p>0.05, * p<0.05, ** p<0.01, *** p< 0.001, **** p<0.0001).

## Results

### Phage display screening and down-selection of lead VHH candidate

In order to identify novel HTT binders, we screened a humanized VHH display library. The workflow combined peptide- and protein-based selection strategies to capture both linear and conformational epitopes using both HTT(Q46) exon 1 fragment as well as N-terminal and polyproline peptides as baits (Figure 1A-B). Following multiple rounds of panning (Supplementary Table 1), 510 unique clone populations were recovered across the screening arms. Sequence analysis revealed broad CDR3 diversity, confirming robust library coverage and a wide range of binding characteristics.

**Figure 1:**
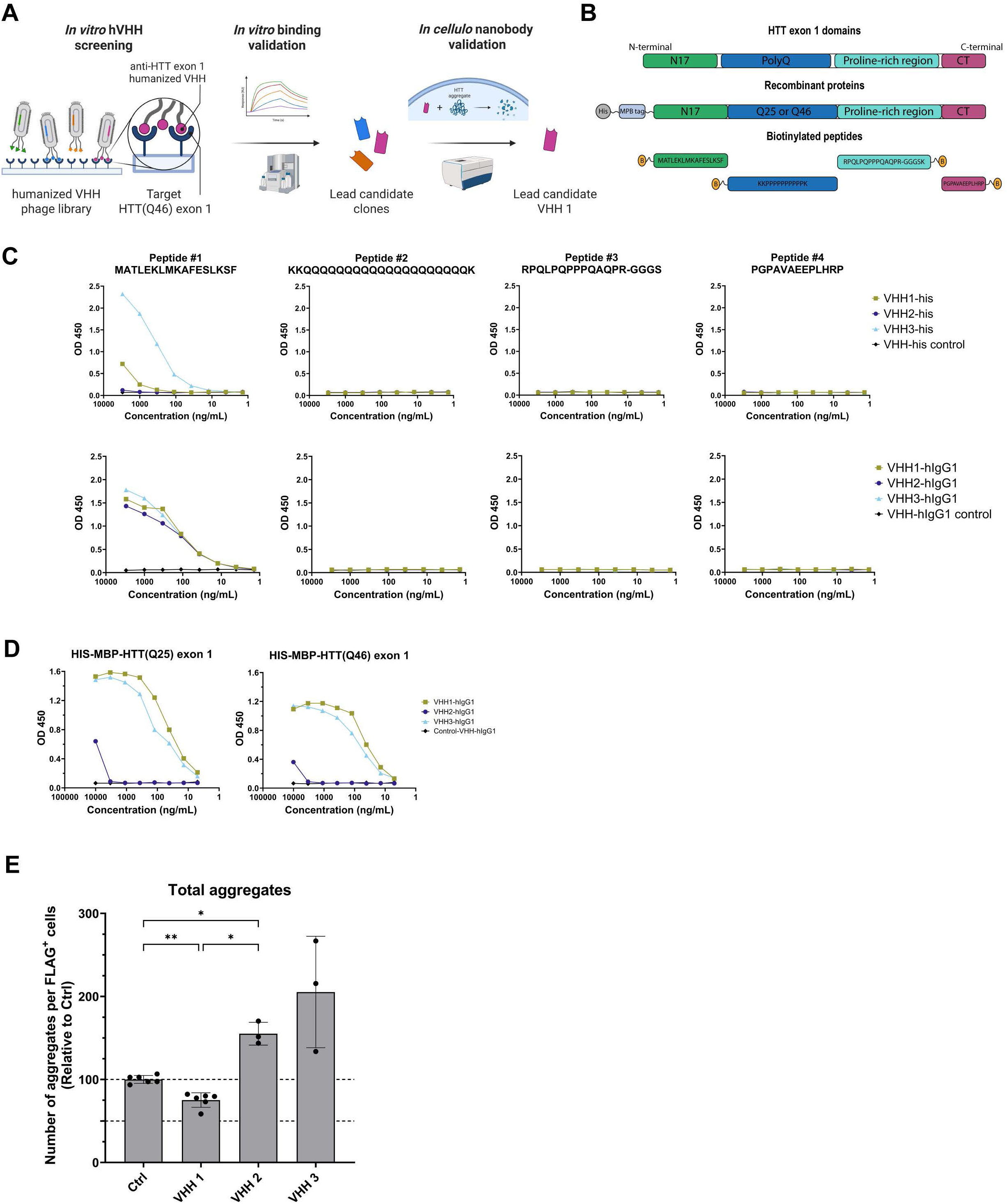
Phage display screening and down-selection of lead VHH candidate. (**A**) Workflow for the discovery and validation of N17-directed intrabodies against HTT exon 1. Intrabodies were selected from a humanized phage display library using recombinant HTT exon 1 (Q46) as the target antigen. Primary screening by AlphaLISA and secondary ranking by surface plasmon resonance (SPR) identified three high-affinity N17 binders (VHH 1, VHH 2, and VHH 3). These lead candidates were expressed as vectorized intrabodies and tested in cell lines for their capacity to reduce intracellular mHTT(Q97) exon 1 aggregates. (**B**) Binding of the purified recombinant VHH-his tagged clones (**Top row**) and the VHH-hIgG1 clones (**Bottom row**) to selected peptides: #1 MATLEKLMKAFESLKSF, #2 KKQQQQQQQQQQQQQQQQQQQQK, #3 RPQLPQPPPQAQPR-GGGS, and #4 PGPAVAEEPLHRP in a peptide based ELISA. (**C**) VHH-hIgG1 clones were examined for binding to directly coated HIS-MBP-HTT exon 1 Q25 and Q46 by ELISA. The reference protein served as the negative control. (**D**) U2OS mHTT(Q97)-IRES-GFPQ16 cells after 72 hours expression of mHTT(Q97) exon 1 and FLAG-tagged intrabody. Number of mHTT(Q97) exon 1 aggregates per transfected FLAG^+^ cell measured by microscopy. Horizontal dotted lines represent 100% and 50% of the control level. Graph represents the mean±SD of three independent experiments (n=3) analyzed by one-way ANOVA (F(0.9916,4.710)=30.31) with Tukey’s multiple comparisons test.

Next, primary screening was done using AlphaLISA. Four sequences covering mHTT exon 1 were used to examine interaction with the remaining clones. In total, 322 VHHs were identified to bind to the N17 and proline-rich region (PRR) peptides, with most showing a strong preference for the N17 region. This finding aligns with previous observations indicating the region’s accessibility and structural stability (De Genst et al., 2015; Gallo et al., 2025). Secondary screening by surface plasmon resonance (SPR) ranked clones based on affinity and specificity. The VHHs were captured using an α-V5 antibody, and binding to recombinant MBP-tagged HTT exon 1 proteins carrying either 25 or 46 glutamines was determined. Seven clones displayed nanomolar affinities, of which three, designated VHH 1, VHH 2, and VHH 3, were the most robust (Supplementary Table 2). These top binders exhibited reproducible kinetics, stable interaction with N17-derived peptides, and no detectable cross-reactivity with unrelated proteins. The selected VHHs were advanced for recombinant production and intracellular validation to assess their functional efficacy.

Epitope mapping and binding analyses by peptide-based ELISA showed that all three VHHs recognize the N17 region of HTT exon 1 fragment in both His-tagged (VHH-His) and IgG1-fused (VHH-IgG1) formats (Figure 1C). However, they display distinct residue dependencies and modification sensitivities. VHH 2 bound the N-terminal portion of N17 and lost binding when residues M1, L4, E5, or L7 of the N17 region were substituted with alanine (Supplementary Table 3). VHH 1 and VHH 3 recognized a more central region centered around F11, with VHH 1 losing binding upon mutation of M8, F11, or E12, and VHH 3 being most dependent on F11 for stable interaction. Post-translational modification (PTM) scanning revealed that VHH 3 maintained binding across all PTMs tested, whereas VHH 1 and 2 were sensitive to S13 or T3 phosphorylation, respectively (Supplementary Table 4).

ELISA was used to determine polyQ length-dependent binding by using recombinant MBP-HTT exon 1 Q25 and Q46 proteins. VHH 1 and VHH 3 bound both antigens with comparable affinities (Figure 1D), confirming that binding is independent of the polyQ length and that the epitope recognition is outside the polyQ tract. VHH 1 displayed the highest apparent affinity and signal intensity, followed by VHH 3, while VHH 2 bound more weakly but remained detectable in both antibody formats.

To determine the effects of the three VHH candidates on untagged mHTT exon 1 fragment aggregation, we transfected plasmids encoding VHH 1, VHH 2, or VHH 3 into human U2OS cells that express mHTT(Q97)-IRES-GFPQ16 upon doxycycline induction. Separated by an IRES, both the mHTT(Q97) exon 1 fragment and GFP-Q16 are controlled by the same promoter but expressed as two separate proteins. This cell line enables us to examine untagged mHTT(Q97) exon 1 aggregation by microscopy using the fluorescent reporter GFP-Q16 that is normally distributed diffusely throughout the cell except when it is sequestered into mHTT aggregates (Geijtenbeek et al., 2022). As negative control, a VHH intrabody targeting influenza virus nucleoprotein 1 (NP1) (Ashour et al., 2015) was included. All VHHs were tagged at the N-terminus with FLAG to allow detection of transfected cells and VHH levels. After 72 hours of mHTT(Q97) exon 1 and VHH expression, aggregate formation was quantified by high-content confocal imaging. Among the tested constructs, only VHH 1 induced a reduction in aggregate counts compared to the VHH control (Figure 1E). Overall, the initial screening and down-selection pipeline identified VHH 1 as a lead candidate that displays specific binding to the N-terminal N17 region of HTT exon 1 and reduces mHTT(Q97) aggregate formation in cells.

### VHH 1 intrabody reduces mHTT aggregation in STHdh cells

Since the striatum is the brain region most severely affected in HD patients, we used mouse striatal cell lines to further investigate the effects of our newly identified VHH 1 on untagged mHTT exon 1 levels. We delivered the intrabodies into mouse striatal STHdh cells that express mHTT(Q97)-IRES-GFPQ16 upon doxycycline induction (Geijtenbeek et al., 2022). Besides the NP1 VHH control, we also included the scFv C4 in our analyses for comparison. Electroporation was used for a high transfection efficiency of more than 80% for all three intrabodies (Supplementary Fig. 1A). After 72 hours of simultaneous expression of mHTT(Q97) and intrabody, we quantified the percentage of intrabody-positive cells with aggregates by automated microscopy. VHH 1 significantly decreased the percentage of cells with aggregates by more than 50% compared to the VHH control, with a similar efficiency as C4 (Fig. 2A). In order to ensure the VHH control did not affect mHTT aggregation by itself, we compared the percentage of cells with aggregates in cells electroporated with different empty plasmid backbones as well as cells that were electroporated without DNA or not electroporated at all. The results showed that the VHH control does not affect mHTT(Q97) aggregation (Supplementary Fig. 1B).

**Figure 2:**
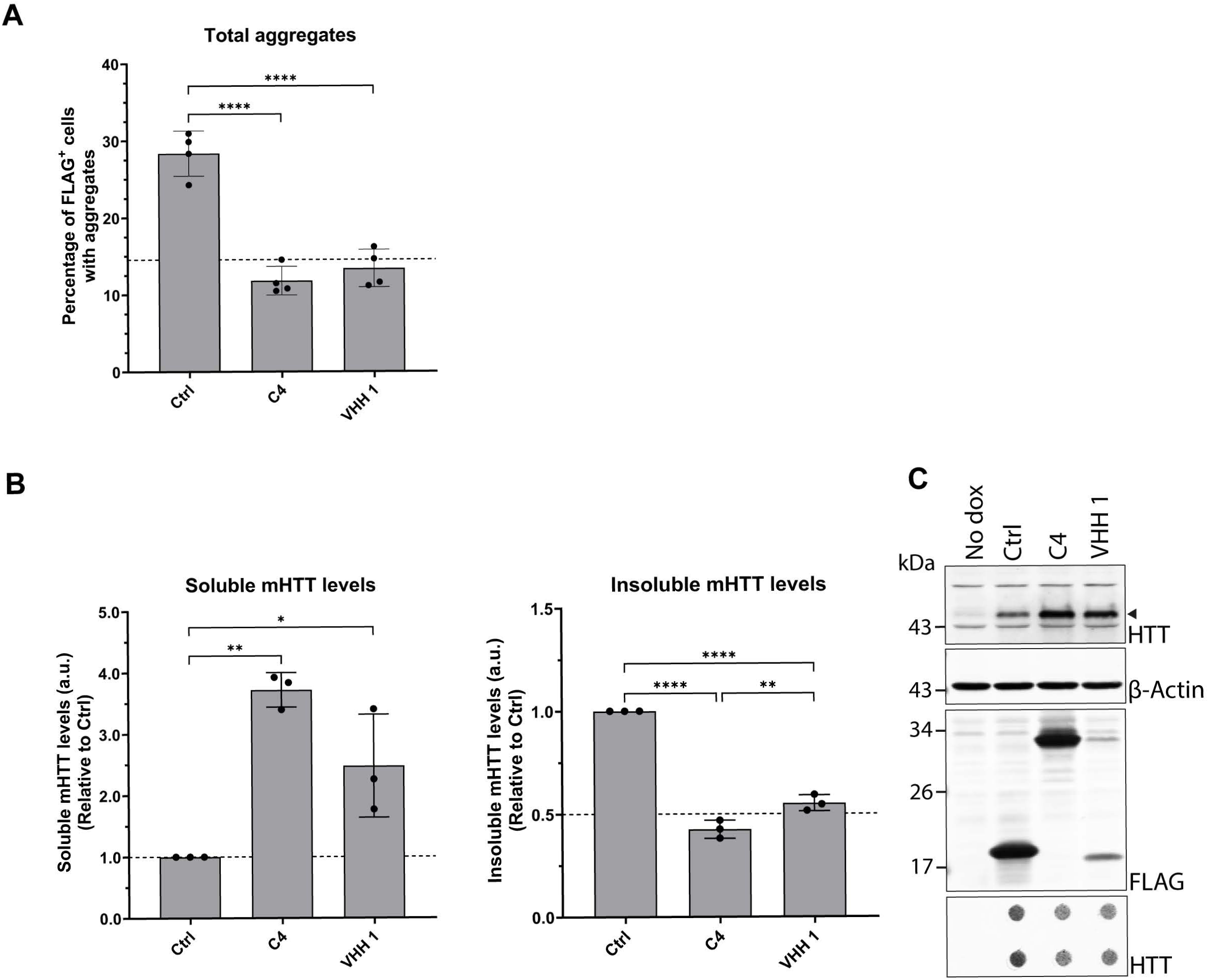
VHH 1 intrabody reduces mHTT aggregation and increases soluble mHTT levels. Soluble and insoluble mHTT(Q97) exon 1 levels in STHdh mHTT(Q97)-IRES-GFPQ16 cells after 72 hours expression of mHTT(Q97) exon 1 and FLAG-tagged intrabody. (**A**) Percentage of transfected FLAG^+^ cells with mHTT(Q97) aggregates measured by microscopy. Horizontal dotted line represents 50% of the control (Ctrl) level (one-way ANOVA, F(2,9)=55.80, n=4). (**B**) Quantification of soluble and insoluble mHTT(Q97) protein levels measured by SDS-PAGE and filter trap assays, respectively, and detected by α-HTT EPR5526 antibody (arrowhead in (**C**)) (one-way ANOVA, Soluble: F(2,6)=21.30, Insoluble: F(2,6)=239.5, n=3). Soluble mHTT(Q97) exon 1 levels were first normalized to β-actin levels before normalizing to the control. Horizontal dotted lines represent 100% (left graph) or 50% (right graph) of the control level. (**C**) Representative blots of the SDS-PAGE blot probed with HTT, actin, and FLAG antibodies as well as the filter trap blot (lower panel) probed with HTT antibody. All graphs represent the mean±SD of at least three independent experiments (n≥3) analyzed by one-way ANOVA with Tukey’s multiple comparisons test.

Consistent with the microscopy observations, both VHH 1 and C4 induced a significant reduction in the level of insoluble mHTT(Q97) exon 1 protein as measured by filter trap assay (Fig. 2B, C lower panel). The decrease in aggregates was accompanied by a concomitant increase in soluble mHTT(Q97) for both VHH 1 and C4 compared to the VHH control as observed by SDS-PAGE analysis (Fig. 2B, C upper panel). These results suggest that similarly to the C4 scFv, VHH 1 can prevent mHTT aggregation by maintaining mHTT in a soluble state.

### Optimizing VHH 1 solubility and expression

To examine the subcellular localization of the intrabodies, cells were stained for HTT and for FLAG to detect the intrabodies prior to imaging by confocal microscopy. The images revealed that all three intrabodies are primarily cytoplasmic (Fig. 3A). However, we also found cells transfected with VHH 1 displaying numerous puncta, indicating solubility issues with the intrabody. Hence, we introduced two point mutations in VHH 1 sequence based on prior studies demonstrating improved stability of single-domain antibodies through specific mutations (Dingus et al., 2022). The resulting optimized intrabody, VHH 1a, presented diffuse cytoplasmic distribution devoid of any puncta, indicating improved solubility compared to the original VHH 1 (Figure 3A).

**Figure 3:**
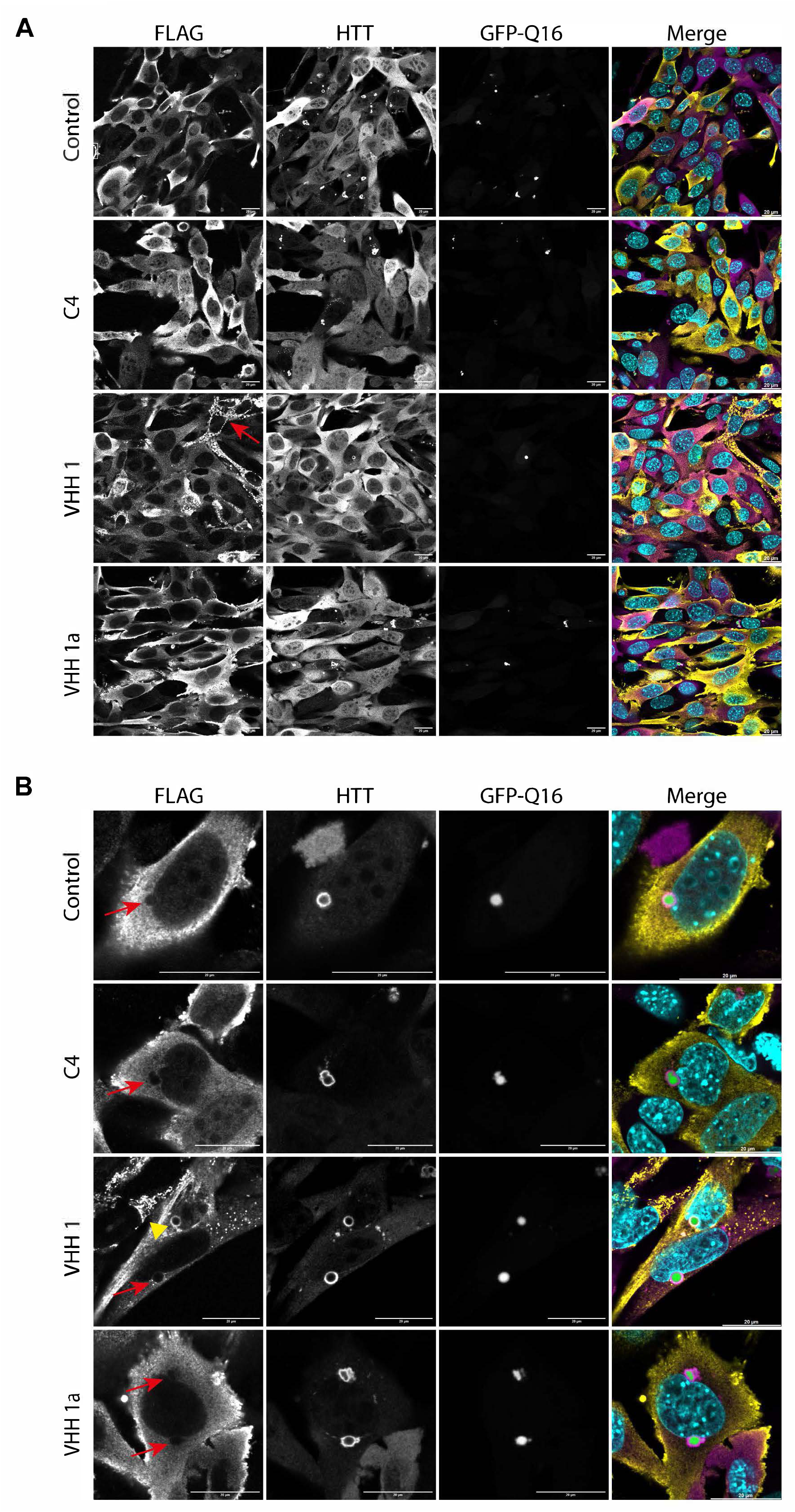
Intrabodies subcellular localization. Representative confocal images of STHdh mHTT(Q97)-IRES-GFPQ16 cells after 72 hours expression of mHTT(Q97) exon 1 and FLAG-tagged intrabody. Cells are stained for FLAG (yellow) and HTT (magenta). Nuclei (cyan) are stained with DAPI. Aggregates are visualized by the recruitment GFP-Q16 (green). Scale bars = 20 µm. (**A**) Intrabodies are primarily located in the cytoplasm. VHH 1 occasionally forms puncta (red arrow). (**B**) Intrabodies are not recruited into aggregates (red arrow), except for VHH 1 that is occasionally co-localized with an aggregates (yellow arrowhead).

Finally, we analyzed whether the intrabodies were recruited to the mHTT aggregates. With the exception of VHH 1, none of the intrabodies were localized to the aggregates (Fig. 3B). Although most cells showed no recruitment of VHH 1 to the aggregates, a few cells did show VHH 1 co-localization with aggregates, potentially linked to its solubility issues. Indeed, this phenotype was never observed with its optimized variant, VHH 1a.

### VHH 1a induces the highest reduction in mHTT aggregates

After confirming that the optimized intrabody VHH 1a displayed comparable binding to recombinant MBP-HTT exon 1 Q25 and Q46 proteins as VHH 1 (Figure 4A), we analyzed the effects of the VHH 1a on soluble and insoluble mHTT(Q97) exon 1 protein levels. STHdh mHTT(Q97)-IRES-GFPQ16 cells were electroporated with the intrabodies and induced with doxycycline before being fixed or harvested after 72 hours. Using automated fluorescence microscopy, VHH 1a showed significantly fewer cells with aggregates compared to the VHH control but also compared to C4 (VHH control: mean 29.12 ± 1.16%; C4: mean 7.78 ± 0.16%; VHH 1a: mean 4.39 ± 0.06%) (Fig. 4B, left panel). The reduction in aggregates observed was similar after performing compartment-specific analysis (Fig. 4B, middle and right panel). Filter trap assay also showed a strong decrease in insoluble mHTT(Q97), with VHH 1a showing the highest efficiency (Fig. 4C, D lower panel). As described earlier for VHH 1 and C4, VHH 1a also increased soluble mHTT(Q97) protein, indicating that the intrabodies prevent mHTT aggregation. Together, these results suggest that VHH 1a is the most efficient at reducing insoluble mHTT(Q97), yet none of the intrabodies are able to lower soluble mHTT(Q97) compared to the VHH control. Of note, VHH 1a was found to have much higher protein levels than VHH 1 on western blot (Fig. 4D, FLAG staining), highlighting the improved solubility following the point mutations.

**Figure 4:**
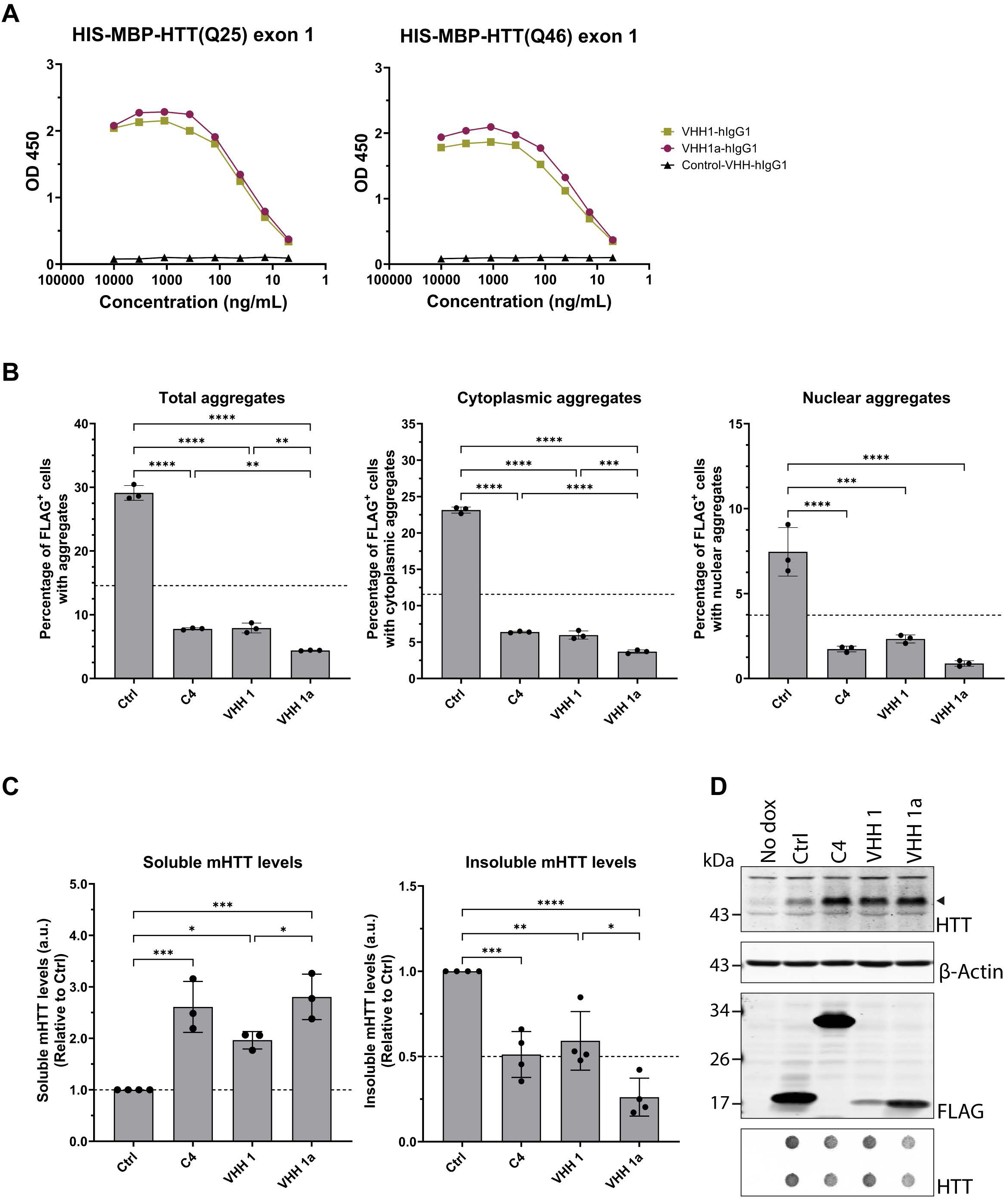
VHH 1a induces the highest reduction in mHTT aggregates. (**A**) VHH-hIgG1 clones were examined for binding to directly coated HIS-MBP-HTT exon 1 Q25 and Q46 by ELISA. VHH 1 and VHH 1a showed comparable binding. (**B-D**) Soluble and insoluble mHTT(Q97) exon 1 levels in STHdh mHTT(Q97)-IRES-GFPQ16 cells after 72 hours expression of mHTT(Q97) exon 1 and FLAG-tagged intrabody. (**B**) Percentage of transfected FLAG^+^ cells with mHTT(Q97) aggregates measured by microscopy (one-way ANOVA, Total aggregates: F(3, 8) = 779.6, Cytoplasmic aggregates: F(3,8)=1710, Nuclear aggregates: F(3,8)=48.95, n=3). Horizontal dotted line represents 50% of the control (Ctrl) level. (**C**) Quantification of soluble and insoluble mHTT(Q97) protein levels measured by SDS-PAGE and filter trap assays, respectively, and detected by α-HTT EPR5526 antibody (arrowhead in (**D**)) (one-way ANOVA, Soluble: F(3,9)=22.60, Insoluble: F(3,12)=25.04, n=4). Soluble mHTT(Q97) exon 1 levels were first normalized to β-actin levels before normalizing to the control. Horizontal dotted lines represent 100% (left graph) or 50% (right graph) of the control level. (**D**) Representative blots of the SDS-PAGE blot probed with HTT, actin, and FLAG antibodies as well as the filter trap blot (lower panel) probed with HTT antibody. All graphs represent the mean±SD of at least three independent experiments (n≥3) analyzed by one-way ANOVA with Tukey’s multiple comparisons test.

In order to verify that the intrabodies do not induce toxicity, a cell count based on nuclear staining was performed by automated microscopy. None of the intrabodies showed a significant change in cell number compared to the VHH control (Supplementary Fig. 2A). Additionally, we performed an MTT assay, which revealed no decrease in cell viability compared to the VHH control (Supplementary Fig. 2B).

### VHH 1a reduces high molecular weight mHTT species but increases small oligomeric ones

As we observed a decrease in insoluble mHTT levels but an increase in soluble mHTT protein, we decided to investigate in more detail the effects of the intrabodies on oligomeric species of mHTT(Q97) exon 1. Due to the resolution of the automated microscope and the pore size of the filter trap membrane, these assays are limited in their detection of mHTT species. As a consequence, some oligomeric species of mHTT may be missed. Therefore, we performed Agarose Gel Electrophoresis for Resolving Aggregates (AGERA) to further investigate the mHTT species composition as this technique has been reported to separate high molecular weight species based on size (Nucifora et al., 2012; Weiss et al., 2008). In order to detect mHTT(Q97) exon 1 levels, blots were stained with either α-HTT S829 or S830 polyclonal antibodies raised against HTT(Q53) exon 1 (Sathasivam et al., 2001). Consistent with the microscopy and filter trap assays, all three intrabodies led to a general decrease in high molecular weight species compared to the VHH control, with VHH 1a showing the strongest reduction (Fig. 5A, B). In addition, a shift in the most abundant mHTT(Q97) species can be observed when analyzing the size at which the signal is the strongest (Fig. 5C). Indeed, both VHH 1 and VHH 1a displayed a shift towards smaller mHTT(Q97) species, whereas C4 showed a shift towards larger mHTT(Q97) species compared to the VHH control. These results hint at a possible difference in the mechanism of action between the VHH intrabodies and C4.

**Figure 5:**
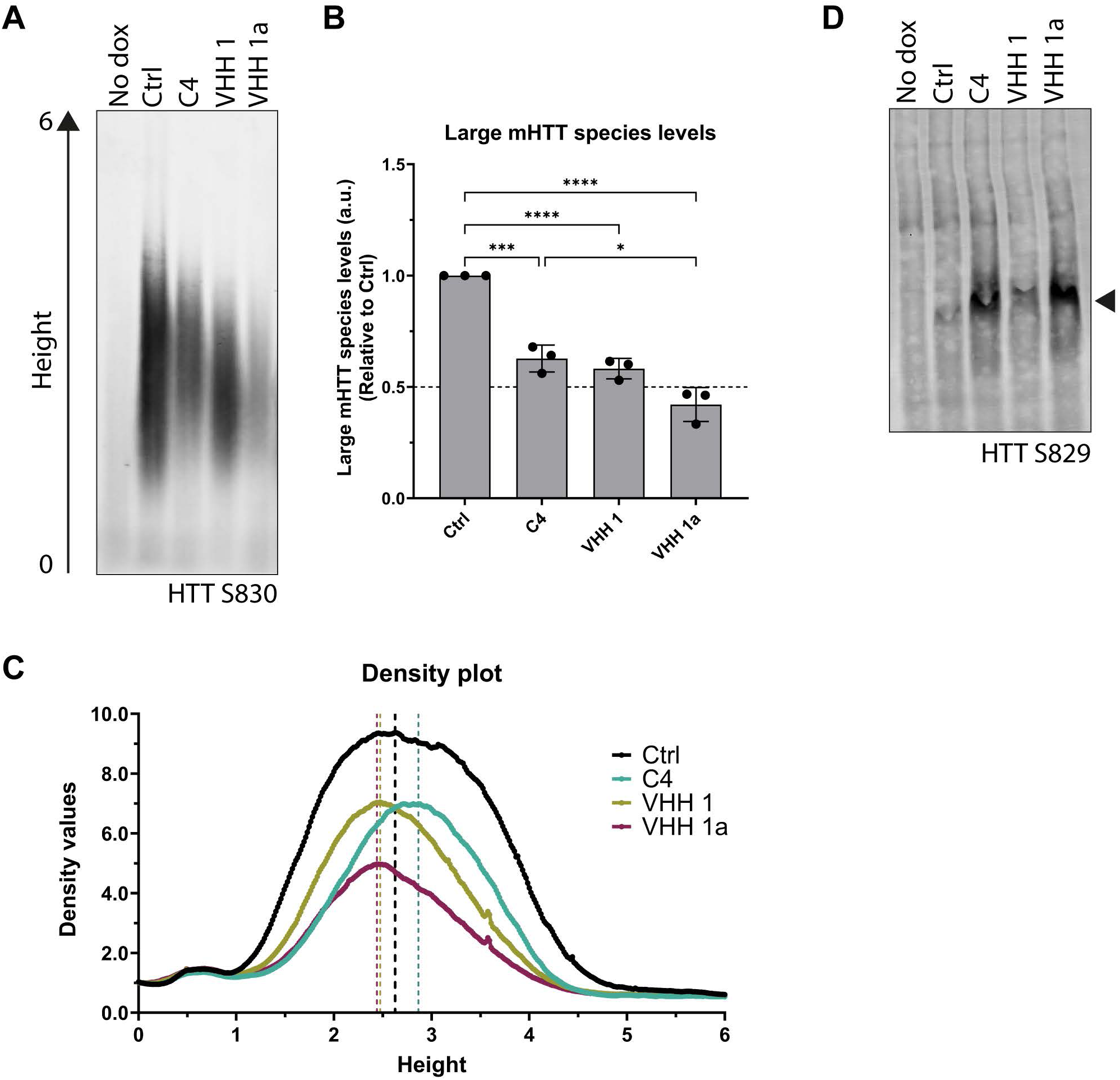
VHH 1a reduces high molecular weight mHTT species but increases small oligomeric ones. Oligomeric mHTT(Q97) levels in STHdh mHTT(Q97)-IRES-GFPQ16 cells after 72 hours expression of mHTT(Q97) exon 1 and FLAG-tagged intrabody. (**A**) Representative blot of Agarose Gel Electrophoresis for Resolving Aggregates (AGERA) probed with α-HTT S830 antibody. (**B**) Quantification of the total smear intensity shows a decrease in high molecular weight mHTT(Q97) species with VHH 1a. The graph represents the mean±SD of three independent experiments analyzed by one-way ANOVA (F(5,12)=58.45, n=3) with Tukey’s multiple comparisons test. The horizontal dotted line represents 50% of the control (Ctrl) level. (**C**) VHH 1 and VHH 1a induce a shift towards smaller mHTT(Q97) species compared to Ctrl on the AGERA as quantified by a density profile plot. The starting and ending distance points are indicated with the left arrow in (**A**). The graph represents the mean of three independent experiments (n=3). Vertical dotted lines represent the distance at which the maximum intensity value is reached for each intrabody. (**D**) Representative blot of a Blue Native PAGE (BN-PAGE) probed with α-HTT S829 antibody showing an increase in small oligomers with VHH 1a compared to Ctrl (arrowhead).

Furthermore, we performed Blue Native Polyacrylamide Gel (BN-PAGE) which is conducted in the absence of SDS and allows for the analysis of smaller oligomeric species (Nucifora et al., 2012). All three intrabodies showed an increase in small oligomeric species compared to the VHH control, particularly VHH 1a and C4 (Fig. 5D). These results align with the increase in soluble mHTT(Q97) species observed by SDS-PAGE analysis. Overall, these results suggest that the intrabodies interfere at the stage of oligomerization to prevent the aggregation of mHTT, thereby reducing the high molecular weight mHTT complexes and increasing the soluble oligomeric mHTT levels.

### VHH 1a specifically targets HTT exon 1, regardless of the polyglutamine stretch length

Since the epitope of the intrabodies is situated in the N17 region of HTT which is common to both full-length protein and exon 1 fragments, we further examined whether the intrabodies could also affect soluble HTT(Q25) exon 1 or full-length HTT(Q7) protein levels. First, STHdh inducible cells that express HTT(Q25)-IRES-GFPQ16 were electroporated and treated with doxycycline, and soluble HTT(Q25) protein levels were analyzed after 72 hours by SDS-PAGE. All intrabodies induced an increase in HTT(Q25) levels compared to the VHH control (Fig. 6A), although the effect was slightly less pronounced than with mHTT(Q97) shown above (Fig. 4B). The effect was much milder with VHH 1, which could be attributed to its lower abundance (Figure 6A, FLAG staining). Next, we investigated the effects of the intrabodies on full-length wild-type HTT(Q7) in wild-type STHdh^Q7/Q7^ or heterozygous STHdh^Q7/Q111^ cells (Trettel et al., 2000) 72 hours after electroporation. None of the intrabodies induced a significant change in full-length HTT(Q7) protein levels (Fig. 6B). These results suggest that the intrabodies only target HTT exon 1 fragment, irrespective of the length of the polyglutamine tract, but not the endogenous full-length protein.

**Figure 6:**
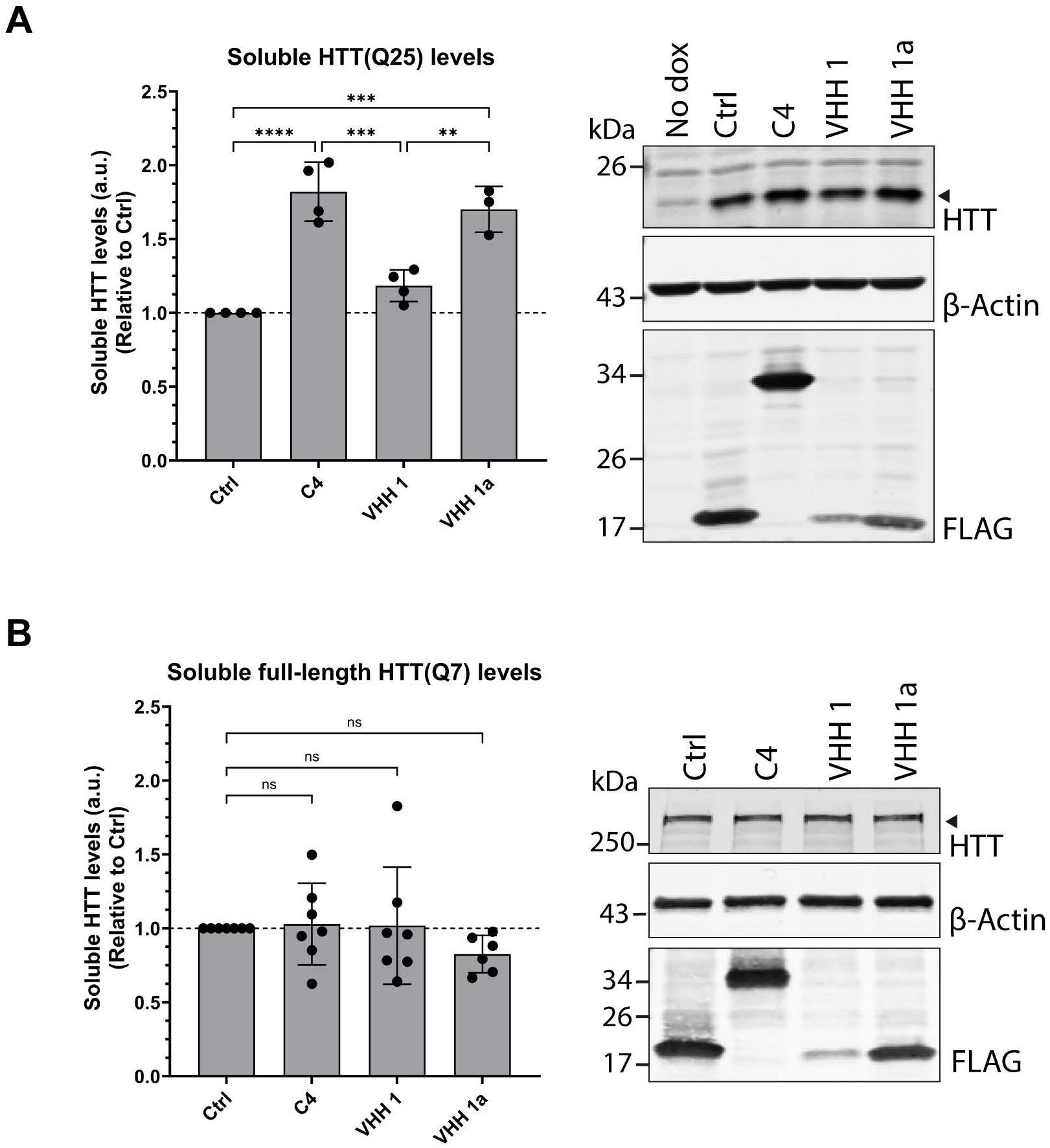
VHH 1a specifically targets HTT exon 1, regardless of the polyglutamine stretch length. **A**) HTT(Q25) exon 1 levels in STHdh HTT(Q25)-IRES-GFPQ16 cells after 72 hours expression of HTT(Q25) exon 1 and FLAG-tagged intrabody as measured by SDS-PAGE and detected by α-HTT EPR5526 antibody (arrowhead). HTT(Q25) exon 1 levels were first normalized to β-actin levels before normalizing to the control (one-way ANOVA, F(3,11)=32.72, n=4). (**B**) Full-length HTT(Q7) levels in STHdh cells after 72 hours expression of FLAG-tagged intrabody as measured by SDS-PAGE and detected by α-HTT EPR5526 antibody (arrowhead) (one-way ANOVA, F(3,23)=0.8829, n=7). All graphs represent the mean±SD of at least three independent experiments (n≥3) analyzed by one-way ANOVA with Tukey’s multiple comparisons test. Horizontal dotted lines represent 100% of the control level.

### VHH 1a prevents further aggregation but does not affect pre-existing aggregates

Since VHH 1a prevents aggregation when expressed simultaneously with mHTT(Q97) exon 1, we investigated whether the intrabodies were also able to remove aggregates already present at the time of intrabody expression. STHdh mHTT(Q97)-IRES-GFPQ16 cells were first exposed to doxycycline for 48 hours to induce mHTT(Q97) exon 1 expression and aggregate formation prior to electroporation with intrabodies (Fig. 7A). Upon electroporation, half of the samples were maintained on doxycycline for continuous mHTT(Q97) expression (+/+ dox samples), whereas the other half were seeded without any additional doxycycline to prevent further mHTT(Q97) expression (+/- dox samples). Cells were then fixed or harvested 72 hours after electroporation. Additionally, cells were harvested after 48 hours of doxycycline just before electroporation.

**Figure 7:**
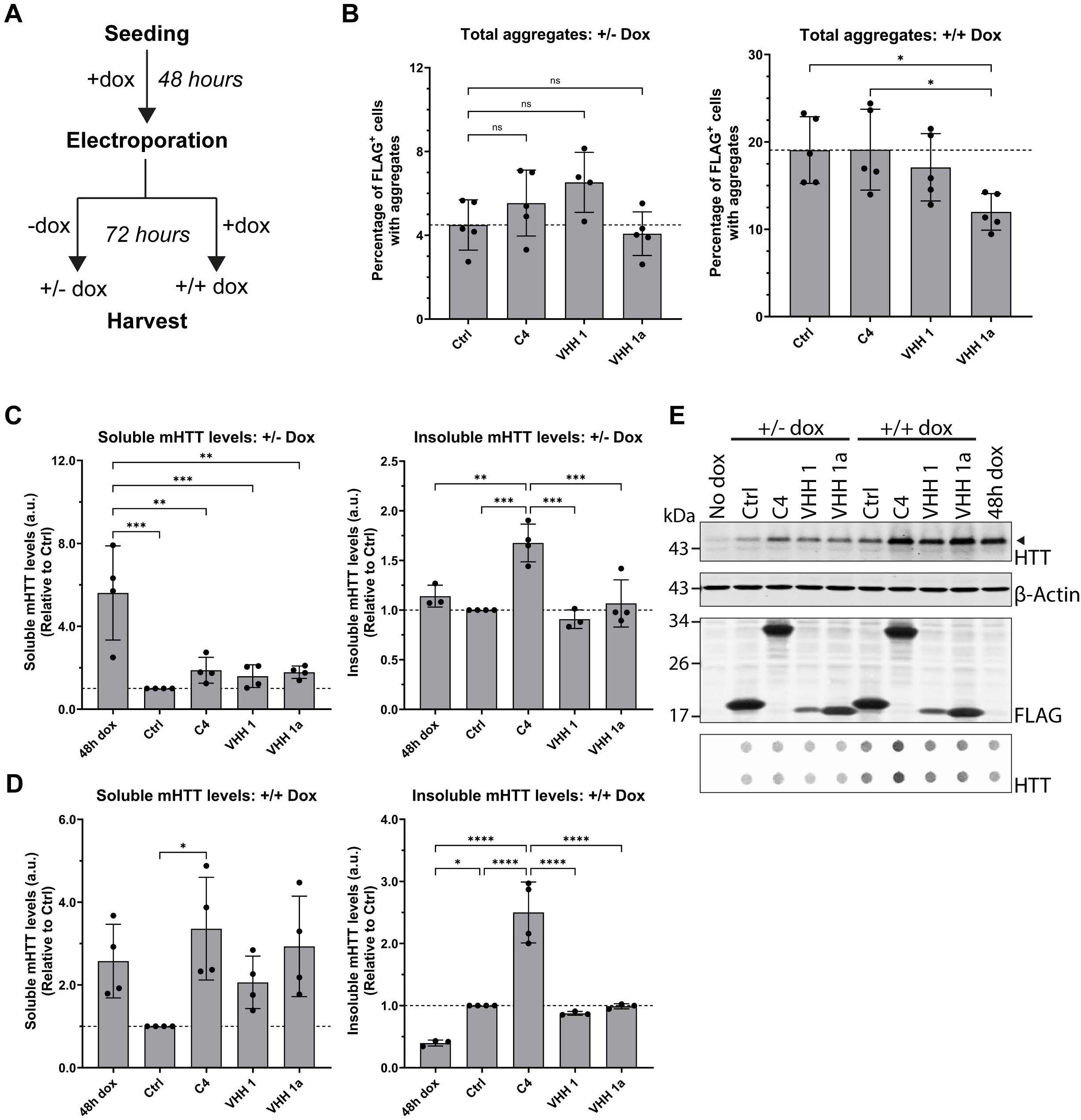
VHH 1a prevents further aggregation without removing pre-formed aggregates. Soluble and insoluble mHTT(Q97) exon 1 levels in STHdh mHTT(Q97)-IRES-GFPQ16 cells after 72 hours expression of mHTT(Q97) exon 1 and FLAG-tagged intrabody in the presence of pre-formed aggregates. (**A**) Experiment timeline schematic. Aggregate formation was induced with doxycycline (dox) for 48 hours prior to electroporation. After electroporation, cells were cultured for 72 hours either with (+/+) or without (+/-) dox before harvest. (**B**) Percentage of transfected FLAG^+^ cells with mHTT(Q97) aggregates measured by microscopy (one-way ANOVA, +/- Dox: F(3,15)=3.095, +/+ Dox: F(3,16)=4.083, n=5). Horizontal dotted lines represent 50% (left graph) or 100% (right graph) of the control level. (**C-D**) Quantification of soluble and insoluble mHTT(Q97) exon 1 levels in +/- dox (**C**) and +/+ dox (**D**) samples measured by SDS-PAGE and filter trap assays, respectively, and detected by α-HTT EPR5526 antibody (arrowhead in (**E**)) (one-way ANOVA, Soluble +/- Dox: F(4,15)=11.40, Insoluble +/- Dox: F(4,13)=14.09, Soluble +/+ Dox: F(4,15)=3.937, Insoluble +/+ Dox: F(4,12)=38.16, n=4). Soluble mHTT(Q97) levels were first normalized to β-actin levels before normalizing to the control. The horizontal dotted line represents 100% of the control level. (**E**) Representative blots of the SDS-PAGE blot probed with HTT, actin, and FLAG antibodies as well as the filter trap blot (lower panel) probed with HTT antibody. All graphs represent the mean±SD of at least three independent experiments (n≥3) analyzed by one-way ANOVA with Tukey’s multiple comparisons test.

Using automated microscopy, we did not observe any significant difference in the percentage of cells with aggregates between the intrabodies and the VHH control when doxycycline was removed prior to intrabody expression, suggesting that none of the intrabodies are able to clear pre-formed aggregates (Fig. 7B, left panel). However, when mHTT(Q97) expression was continued throughout the experiment, VHH 1a displayed significantly fewer cells with aggregates compared to the VHH control, whereas C4 showed similar aggregation level as the control (Fig. 7B, right panel). Instead, VHH 1a maintained the aggregation level to that of the 48-hour doxycycline samples (Supplementary Fig. 3A, +/+dox). Hence, these results suggest that only VHH 1a prevents the formation of additional large aggregates by newly synthesized HTT exon 1 when aggregates are already present in the cells.

When analyzing the levels of insoluble mHTT(Q97) protein by filter trap assay, C4 showed a significant increase in insoluble species whether doxycycline was removed (Fig. 7C, E lower panel) or maintained throughout the experiment (Fig. 7D, E lower panel). When doxycycline was maintained, C4 also significantly increased soluble mHTT(Q97) levels. On the other hand, VHH 1 and VHH 1a maintained the levels of insoluble mHTT(Q97) to that of the VHH control regardless of the doxycycline treatment. The discrepancy between the microscopy and biochemistry results may arise from smaller insoluble species that are too small to be detectable as inclusion bodies by microscopy but large and insoluble enough to be captured by the filter trap membrane. Also, increased sizes of the larger aggregates would not be detected by microscopy analysis but would increase the total signal detected on filter trap membrane. Lastly, the increase in soluble mHTT upon intrabody expression was observed only with continuous mHTT expression (Fig. 7D) but not when doxycycline was removed (Fig. 7C), likely due to the lack of new mHTT production. Overall, these results indicate that VHH 1a prevents further formation of large mHTT aggregates even when aggregates are already present. In contrast, C4 still allows the generation of new aggregates and even exacerbates the formation of insoluble mHTT species on filter trap.

## Discussion

Huntington’s disease is characterized by the accumulation and aggregation of mHTT protein fragments with an extended polyQ tract. An appealing approach to delay or prevent HD onset involves promoting the degradation of mHTT protein. Intrabodies have recently garnered a lot of attention for their therapeutic potential with their ability to specifically target intracellular pathological proteins such as proteins involved in neurological disorders including α-synuclein, tau, and HTT (Messer & Butler, 2020).

The first intrabody developed against HTT was a scFv (C4) targeting the N-terminus of HTT that has been extensively shown to reduce mHTT aggregation and toxicity in various models (Butler & Messer, 2011; Kvam et al., 2009; Lecerf et al., 2001; Miller et al., 2005; Murphy & Messer, 2004; Wolfgang et al., 2005). Although promising, the protective effects of C4 diminished with time in transgenic mice (Snyder-Keller et al., 2010), and *Drosophila* flies still died earlier compared to healthy controls (Wolfgang et al., 2005). Since the development of C4, additional intrabodies targeting different regions of HTT have been published with variable efficacy in reducing HTT aggregates (Denis et al., 2019; Jurcau et al., 2024; Messer & Butler, 2020). Yet, an optimal binder with high and sustainable *in vivo* efficacy and good safety profile remains to be found.

In order to identify new binders, we screened a humanized VHH page-display library for VHH intrabodies able to bind mHTT exon 1 protein, which resulted in the identification of three promising binders specific for the N17 region of HTT. Here, the N17 region is an interesting sequence as it is a key domain that regulates HTT physiological function but also modulates mHTT exon 1 aggregation and toxicity (Arndt et al., 2015; Greiner & Yang, 2011; Vieweg et al., 2021). Although N-terminal binders risk cross-reactivity with the wild-type protein, polyQ-directed binders lack specificity and have even been shown to exacerbate aggregation and toxicity (Khoshnan et al., 2002; Kvam et al., 2009; Legleiter et al., 2009). We investigated the effects of the intrabodies on HTT aggregation using cell lines expressing untagged mHTT exon 1 as tags can substantially alter HTT aggregation and degradation dynamics (Pandey et al., 2024; Riguet et al., 2021; Wagner et al., 2018). VHH 1 was the only intrabody that reduced mHTT aggregates in human U2OS cells, whereas the strongest binder, VHH 3, led to an increase in aggregation. This could be due to conformational differences or to the fact that high affinity may instead promote nucleation and protein aggregation. These results highlight the need to assess the effects of a binder on phenotype in cellular models as a higher *in vitro* affinity does not equate to optimal efficacy.

After optimization of VHH 1 into VHH 1a, we showed that our intrabody was the most efficient at reducing insoluble mHTT exon 1 species without affecting the full-length wild-type protein levels in mouse striatal cells, indicating specificity for the toxic fragment. Interestingly, although both VHH 1a and C4 decrease the overall levels of high molecular weight species, VHH 1a induces a shift towards smaller species, whereas C4 induces a shift towards higher molecular weight species, suggesting that the VHH intrabody is more efficient at inhibiting aggregate growth. Yet, both VHH 1a and C4 increase the level of small oligomeric complexes compared to the VHH control. Hence, the intrabodies appear to act on the small soluble oligomeric mHTT species in order to prevent the transition into large aggregated species.

Additional differences between VHH 1a and C4 were observed when studying the clearance of already-formed aggregates. While neither VHH 1a nor C4 were able to reduce the level of pre-existing aggregates, only VHH 1a was able to prevent the continued formation of new large aggregates. The reduction of insoluble mHTT complexes by VHH 1a compared to the control was observed by microscopy but not by filter trap assay, which suggests that VHH 1a prevents small insoluble mHTT species from being sequestered into inclusion bodies. In contrast, C4 increases the generation of insoluble mHTT complexes and cannot prevent the sequestration of mHTT species into larger complexes when aggregates already exist (Figure 8). Overall, these results highlight a significant differences in the mechanism of action and perhaps even recognition of different mHTT species between VHH 1a and C4. These findings also emphasize the need to assess the intrabody efficacy not only as prophylactic but also as therapeutic agent in order to determine the best approach when it comes to treating patients that may already present some pathology when receiving treatment.

**Figure 8:**
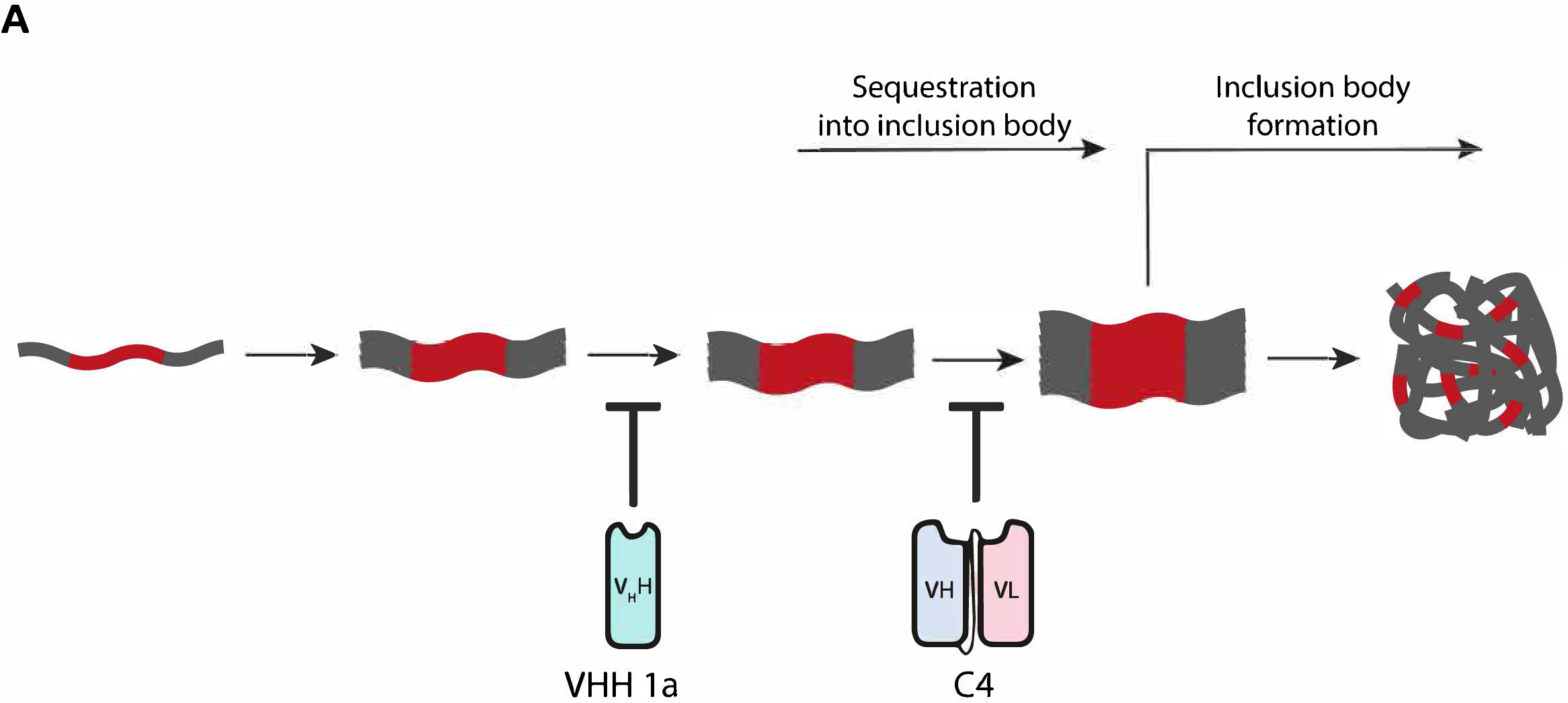
Model for the mechanism of action of the intrabodies on mHTT aggregation. Diagram explaining the proposed mechanism of action of VHH 1a and C4. Both intrabodies prevent the initiation of inclusion body formation. VHH 1a works on different mHTT species than C4 and thereby can prevent the sequestration of newly synthesized mHTT species into existing inclusion bodies. In contrast, if mHTT has already started accumulating, C4 exacerbates the aggregation phenotype by stabilizing a sequestration-prone mHTT species.

## Acknowledgements

We would like to thank Prof. Dr. Gillian Bates (University College London) for kindly providing the S829 and S830 antibodies. Figure 1A was created in BioRender. This work was funded by VectorY Therapeutics B.V, Amsterdam, The Netherlands.

## Notes

### Competing Interest Statement

Eric A.J. Reits reports financial support, administrative support, equipment, drugs, or supplies, statistical analysis, and writing assistance were provided by VectorY Therapeutics BV. Eric A.J. Reits reports a relationship with VectorY Therapeutics BV that includes: funding grants and non-financial support. Eric A.J. Reits has patent pending to VectorY Therapeutics BV. If there are other authors, they declare that they have no known competing financial interests or personal relationships that could have appeared to influence the work reported in this paper.

