## Supplementary Figures for "Development of a novel VHH intrabody targeting the N17 region of huntingtin exon 1 protein that prevents inclusion body formation"

*Supplementary Figure 1*: **Control VHH does not affect mHTT aggregation**. Aggregate level in STHdh mHTT(Q97)-IRES-GFPQ16 cells after 72 hours expression of mHTT(Q97) exon 1 and FLAG-tagged intrabody or control constructs. (**A**) Percentage of transfected $FLAG^+$ cells measured by microscopy is above 70% for all constructs. The graph represents the mean±SD of four independent experiments (n=4). (**B**) Percentage of cells with mHTT(Q97) aggregates is not altered by the VHH control compared to empty backbone constructs (pcDNA; pUC19) and cells without DNA transfection (Electroporation; No electroporation). The graph represents the mean±SD analyzed by one-way ANOVA ($F(4,10)=1.578$, $n \geq 1$) with Tukey's multiple comparisons test.

*Supplementary Figure 2*: **Intrabodies do not induce cytotoxicity**. Cell viability assays in STHdh mHTT(Q97)-IRES-GFPQ16 cells after 72 hours expression of mHTT(Q97) exon 1 and FLAG-tagged intrabody. (**A**) No significant change in cell number per well quantified by Hoechst-stained nuclei count by microscopy (one-way ANOVA, $F(3,8)=0.2524$, n=3). (**B**) No significant differences in cell viability measured by MTT assay (one-way ANOVA, $F(3,8)=1.934$ n=3). All graphs represent the mean±SD of three independent experiments (n=3) analyzed by one-way ANOVA with Tukey's multiple comparisons test. The horizontal dotted line represents 100% of the control level.

*Supplementary Figure 3*: **Total amount of cell with aggregates**. (**A**) Aggregate level in STHdh mHTT(Q97)-IRES-GFPQ16 cells after 72 hours expression of mHTT(Q97) exon 1 and FLAG-tagged intrabody in the presence of pre-formed aggregates. Percentage of total cells with mHTT(Q97) aggregates measured by microscopy either before electroporation (48h sample) or after electroporation with (+/+) or without (+/-) doxycycline maintained. The graph represents the mean±SD analyzed by one-way ANOVA ($F(8,32)=29.44$, n=5) with Tukey's multiple comparisons test. Significant difference is indicated by the compact letter display where identical letters indicate non-significant difference between two groups.

Supplementary Figure 1

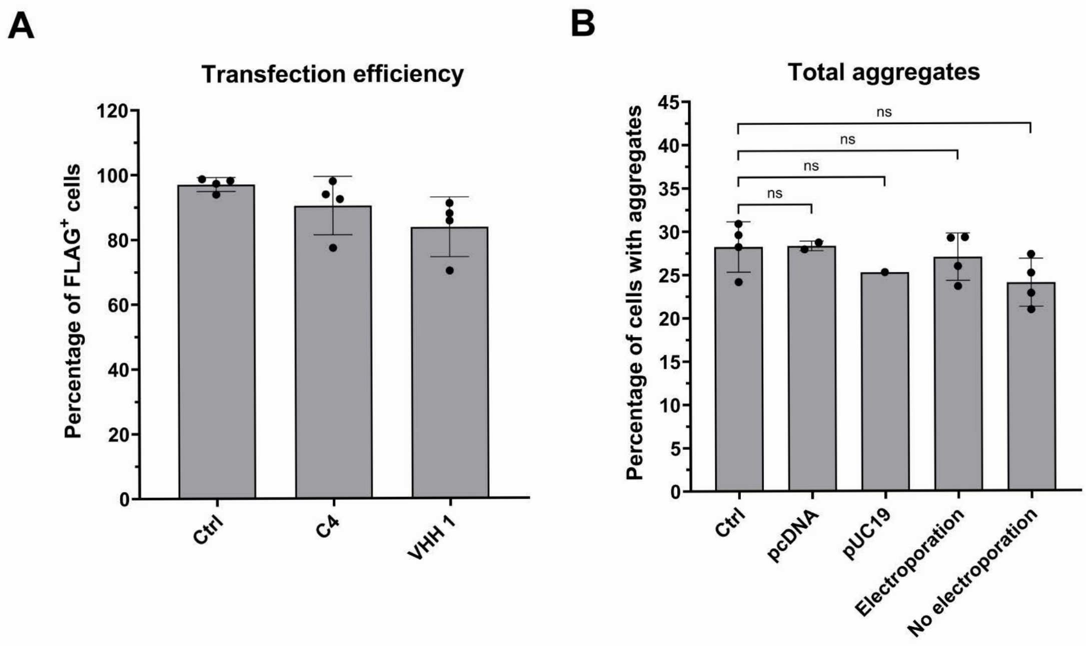

Supplementary Figure 2

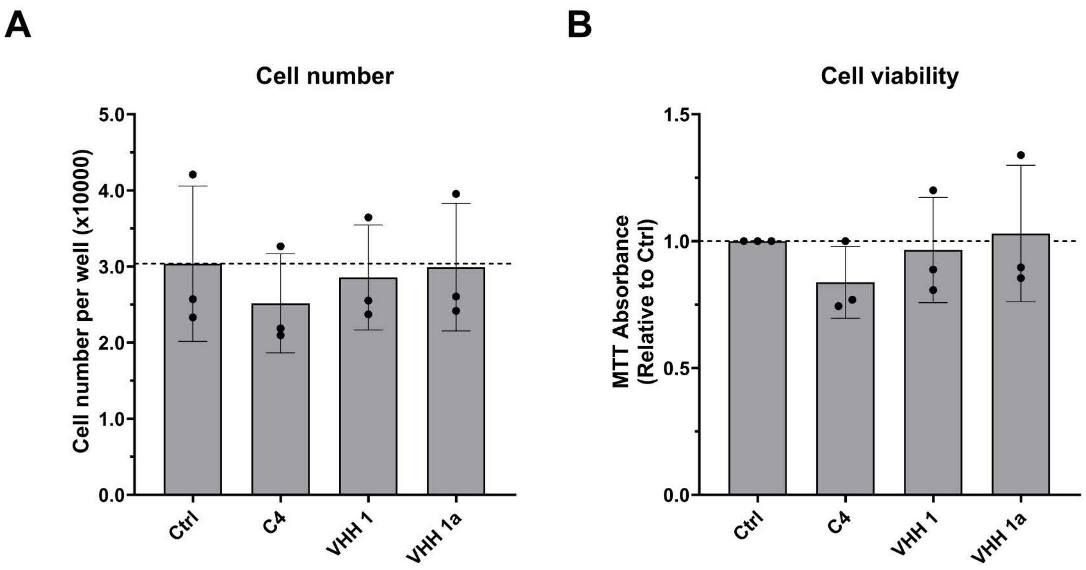

Supplementary Figure 3

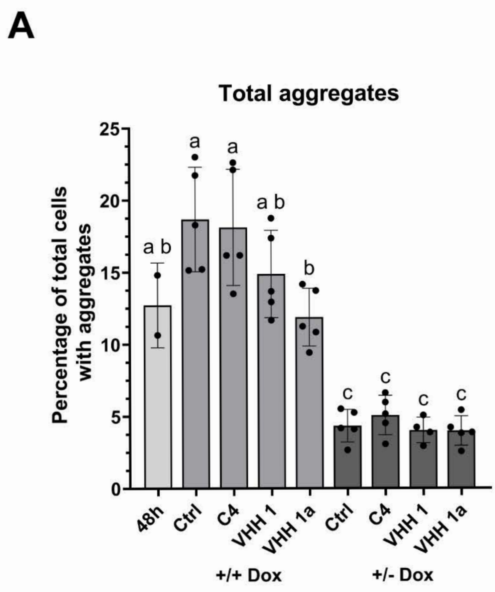

Uncropped and annotated western blots

Figure 2C - Uncropped original blots

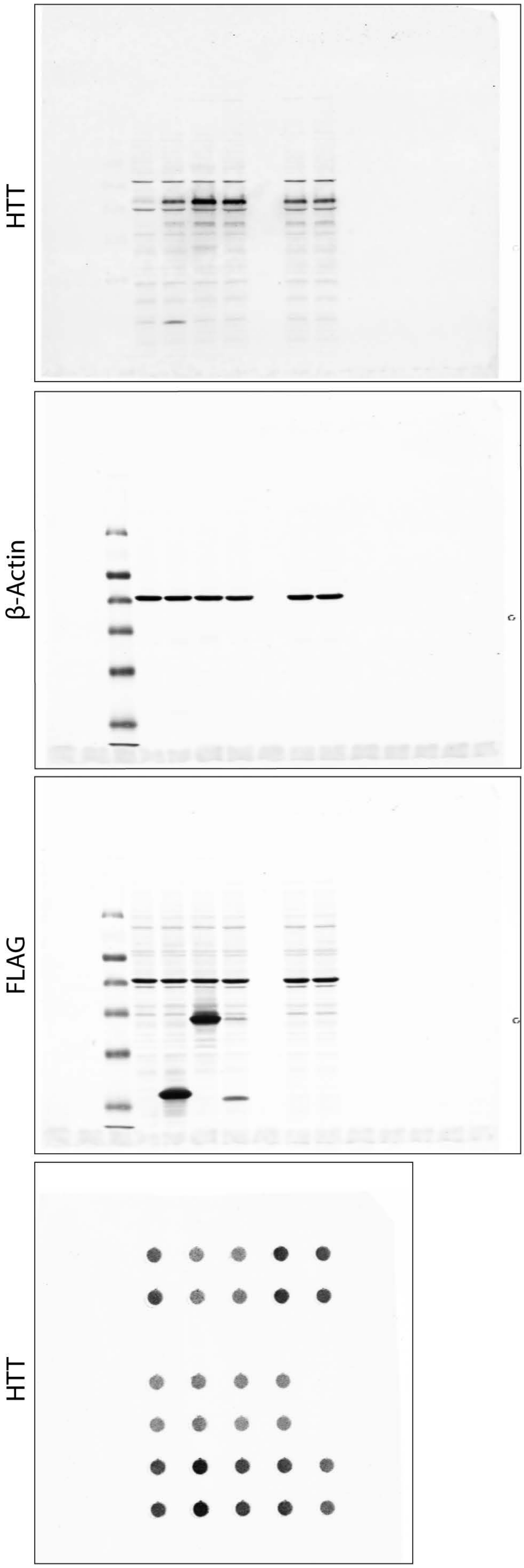


Figure 2C - Annotated blots

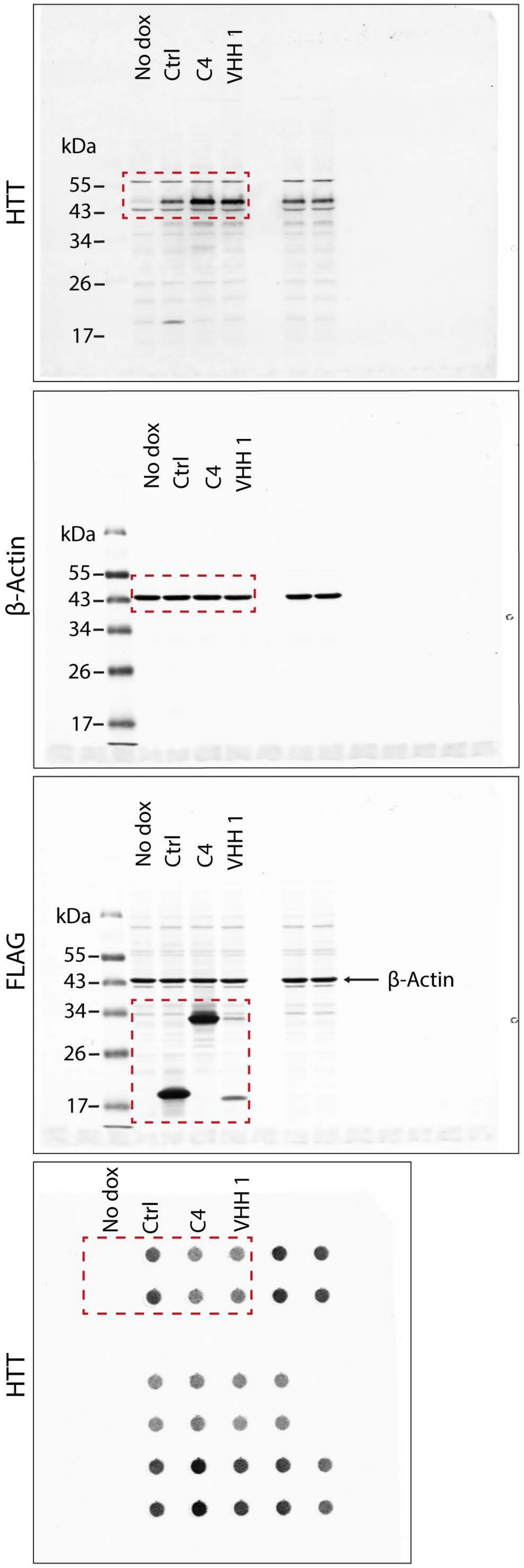

Uncropped and annotated western blots

Figure 4D - Uncropped original blots

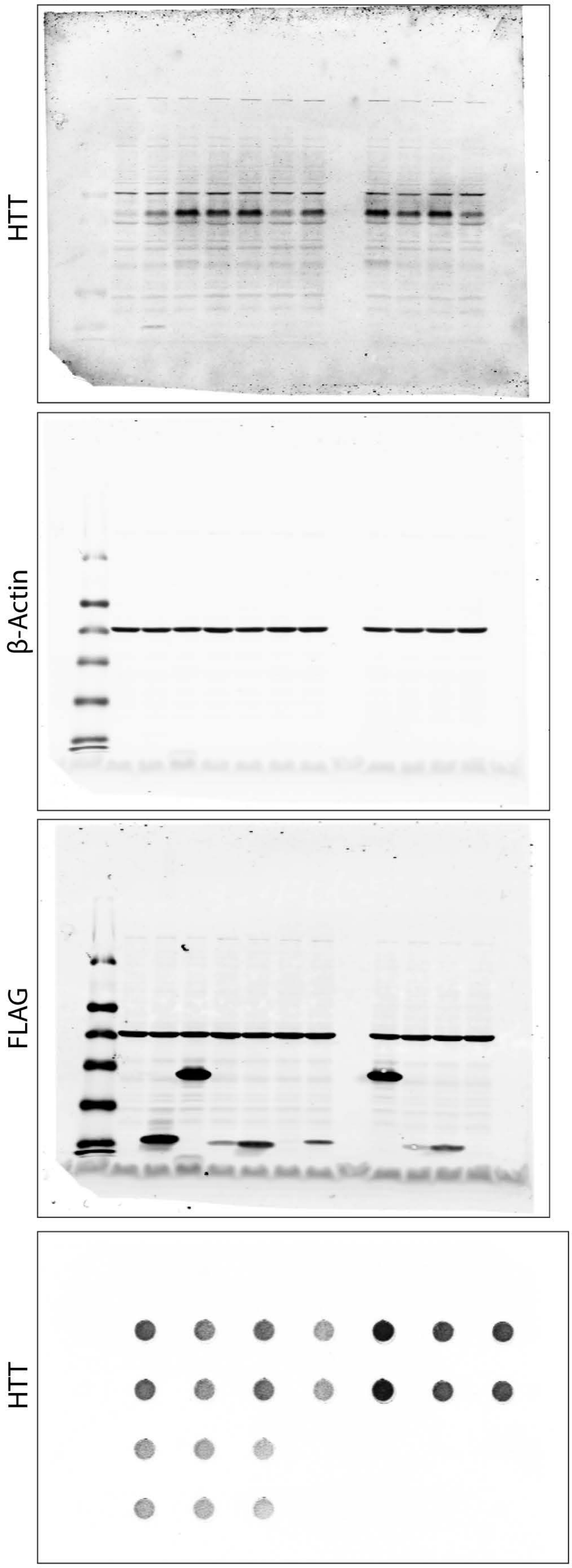


Figure 4D - Annotated blots

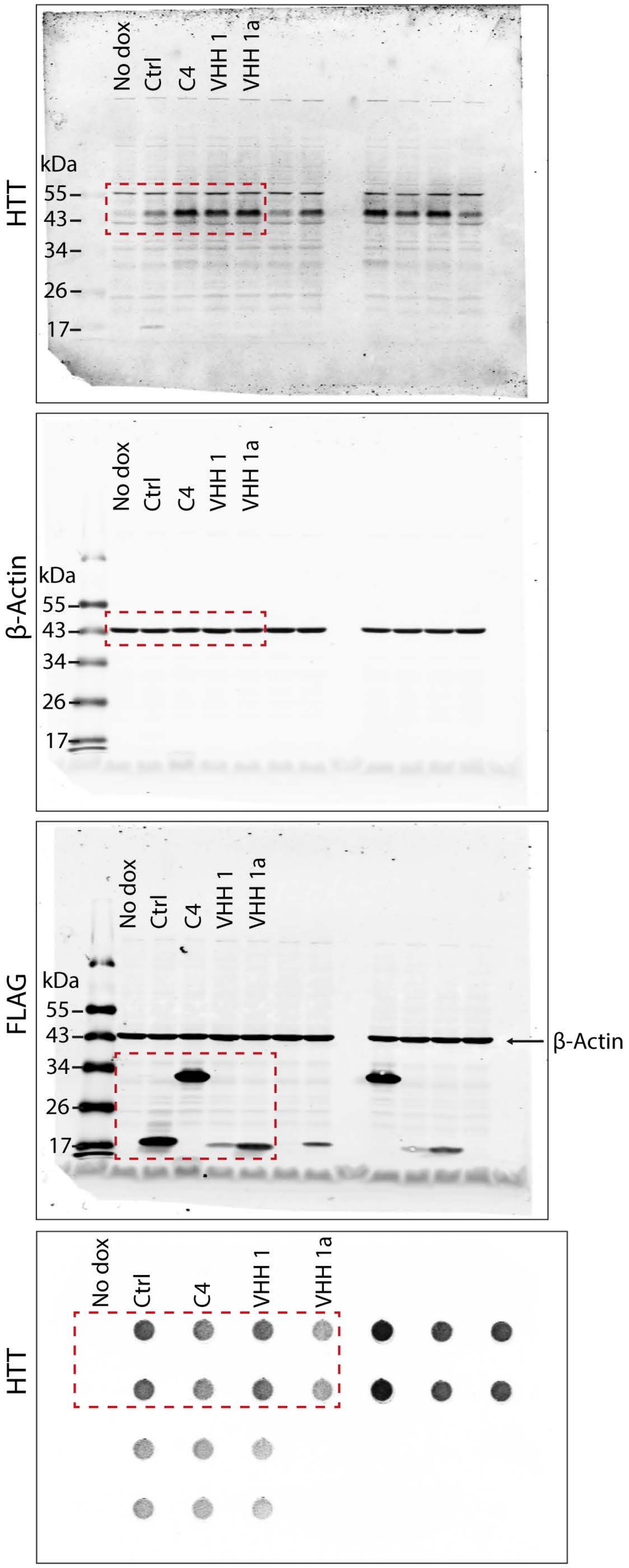

Uncropped and annotated western blots

Figure 5A - Uncropped original blots

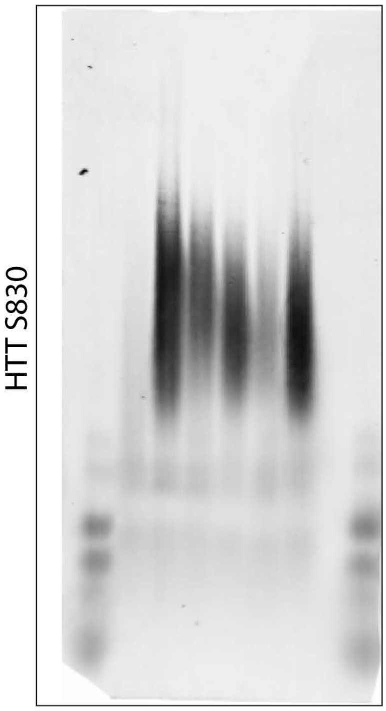


Figure 5A - Annotated blots

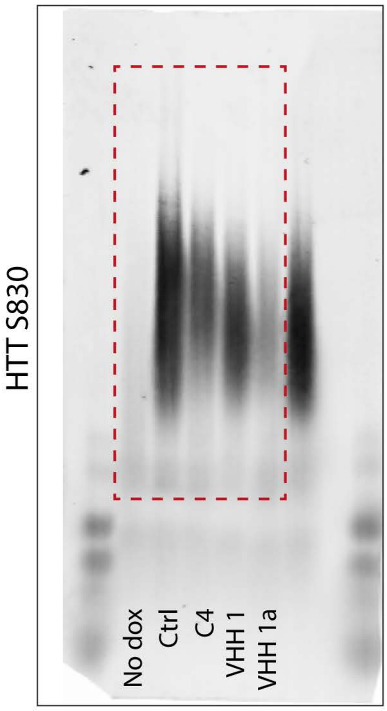


Figure 5D - Uncropped

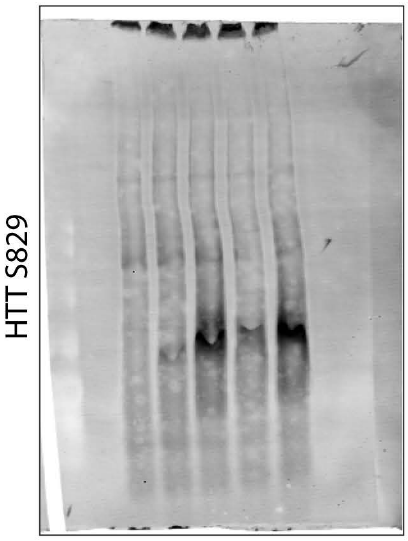


Figure 5D - Annotated

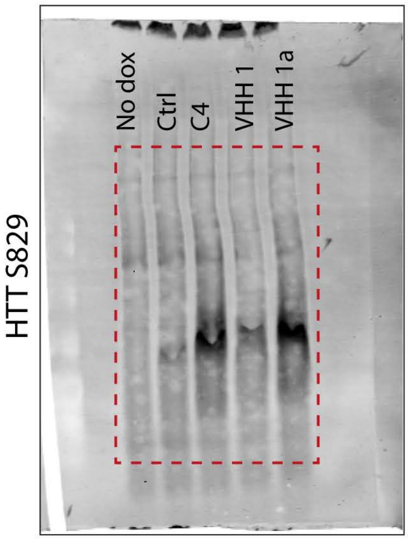

Uncropped and annotated western blots

Figure 6A - Uncropped

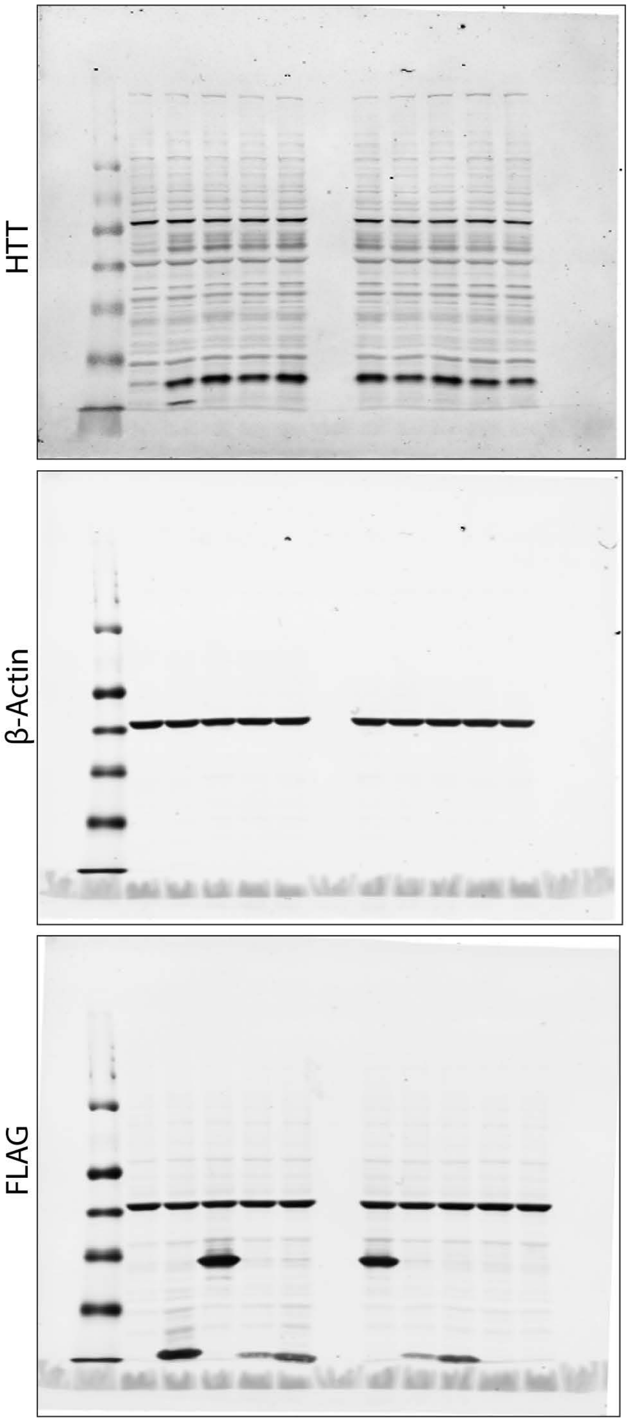


Figure 6A - Annotated blots

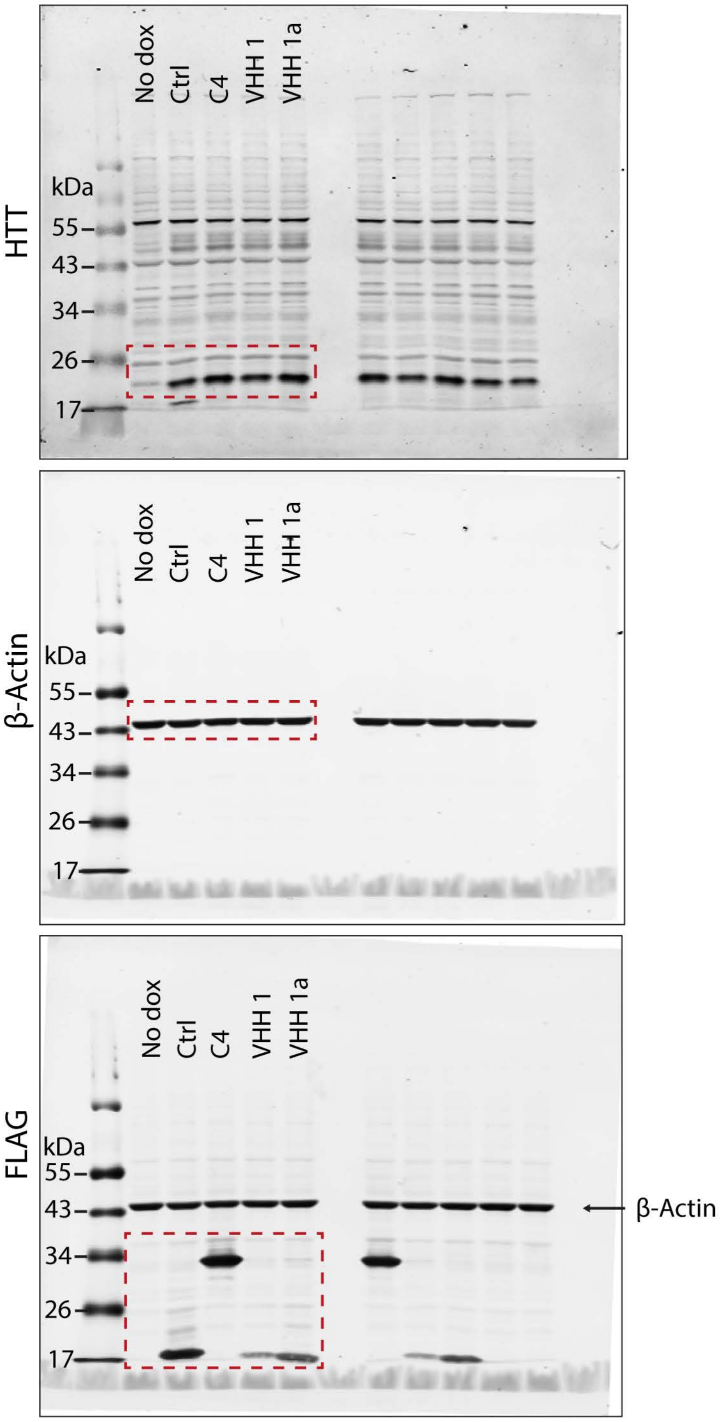

Uncropped and annotated western blots

Figure 6B - Uncropped original blots

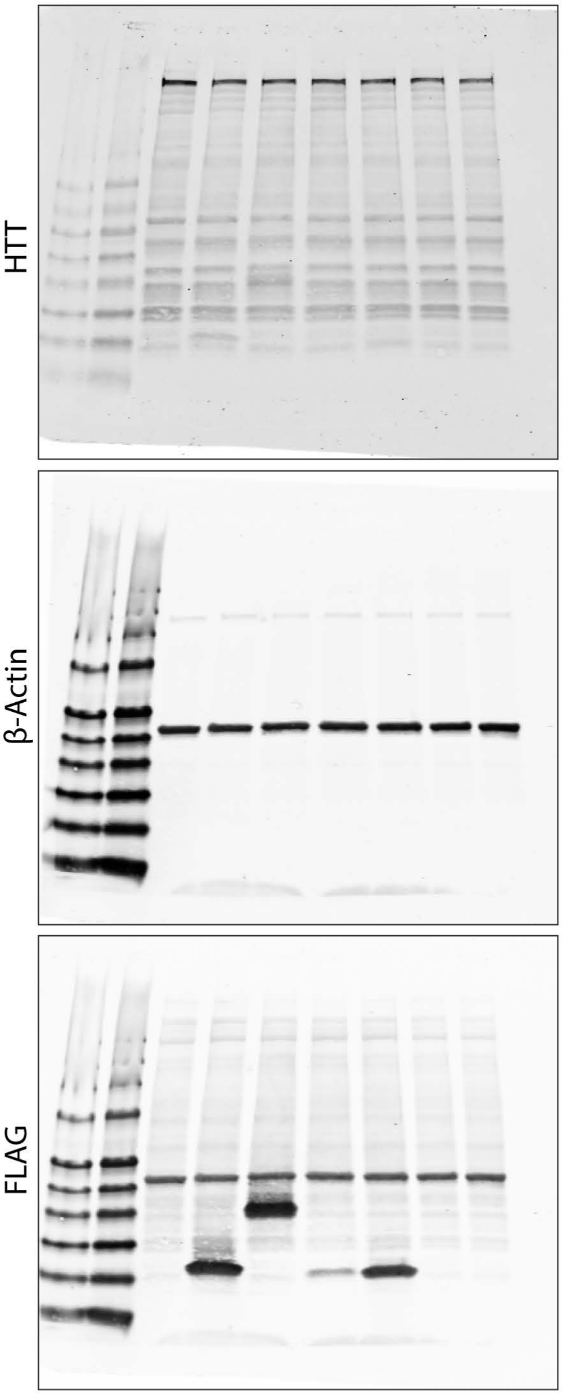


Figure 6B - Annotated blots

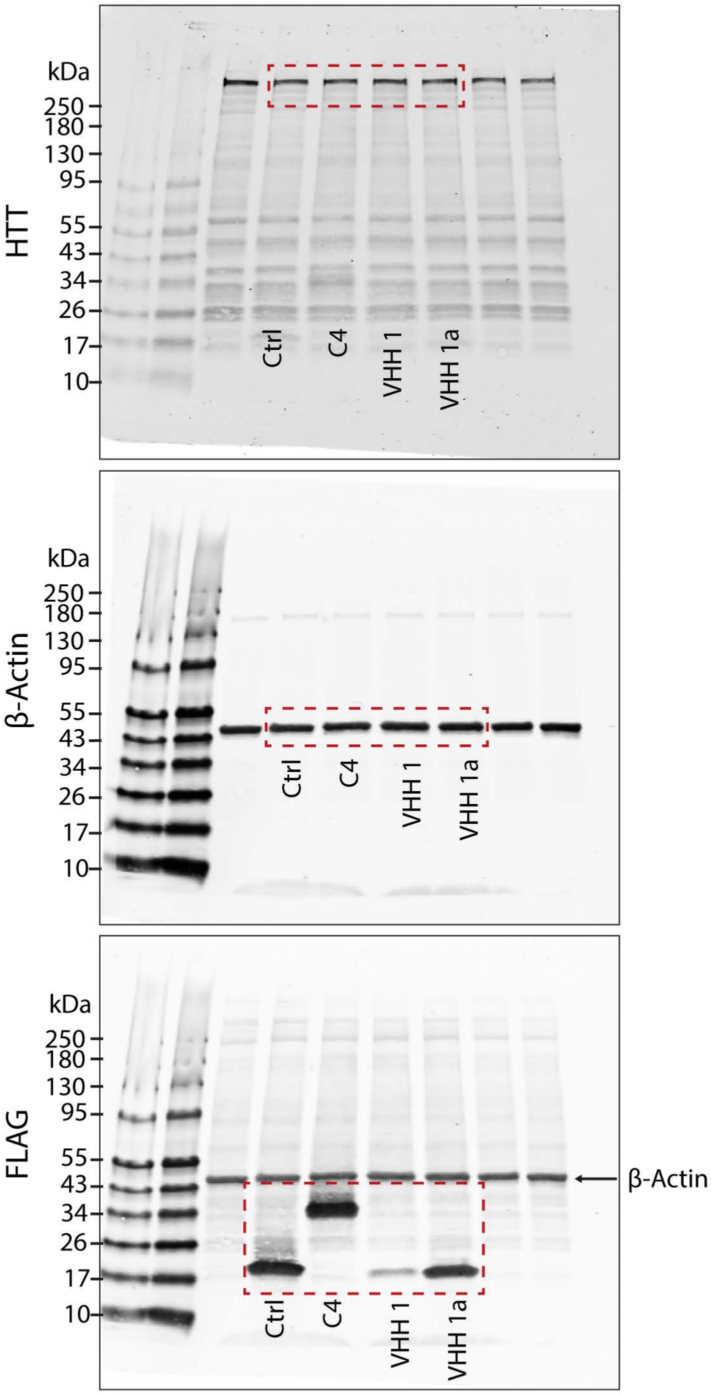

Uncropped and annotated western blots

Figure 7E - Uncropped original blots

HTT

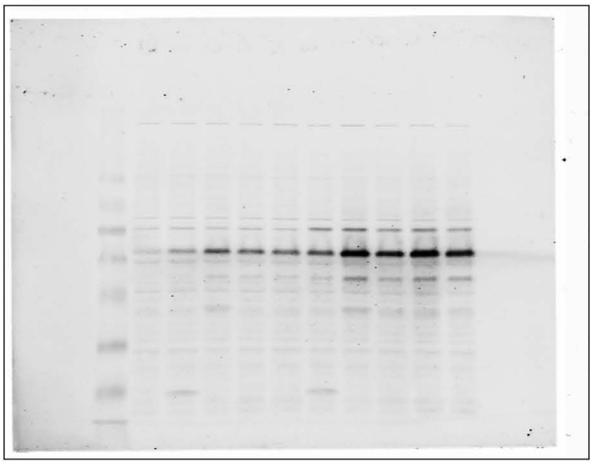

β-Actin

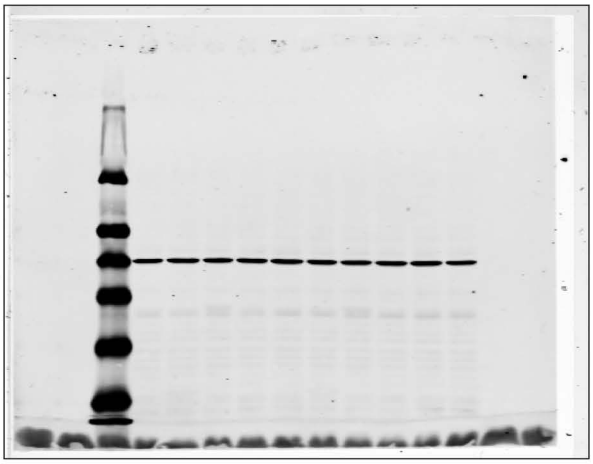

FLAG

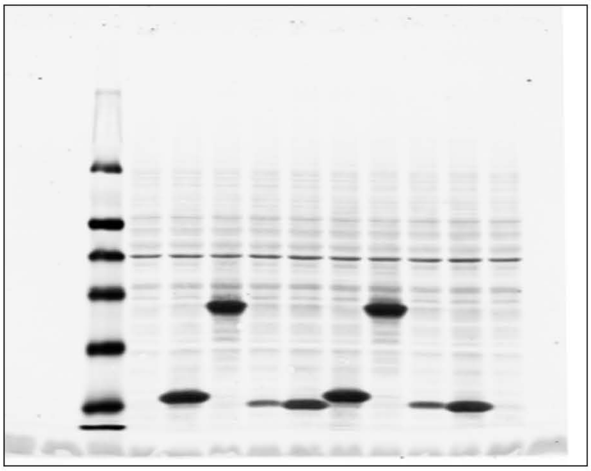

HTT

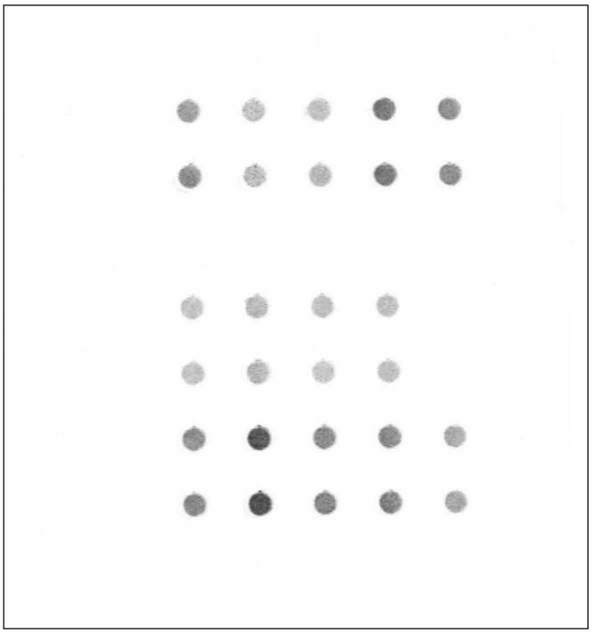

Figure 7E - Annotated blots

HTT

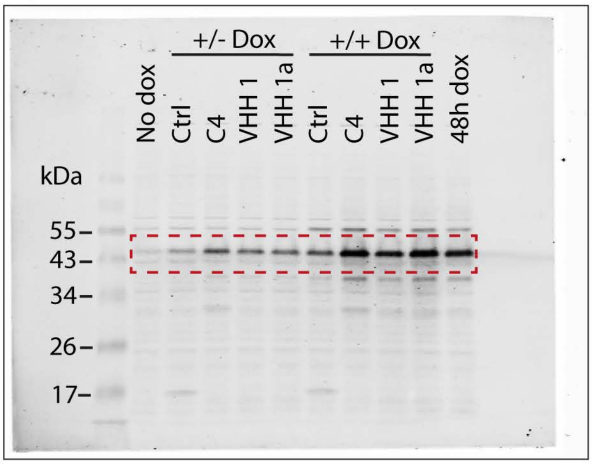


β-Actin

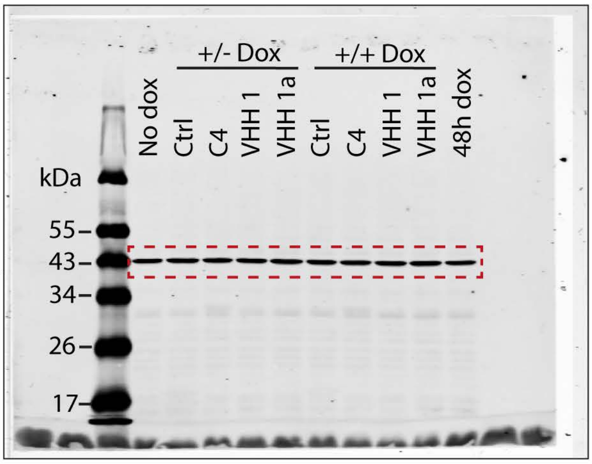


FLAG

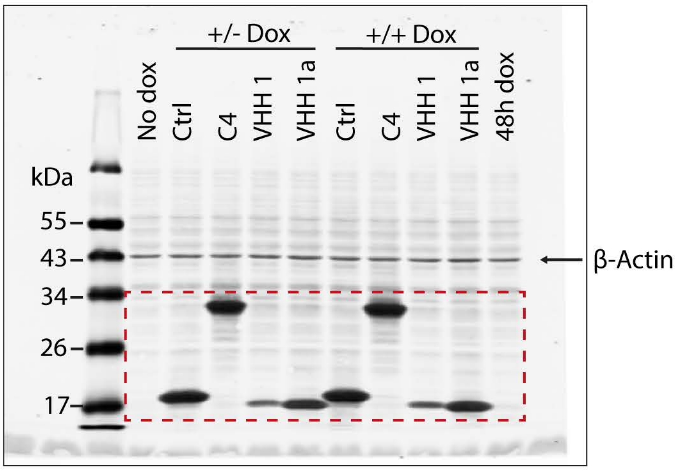


HTT

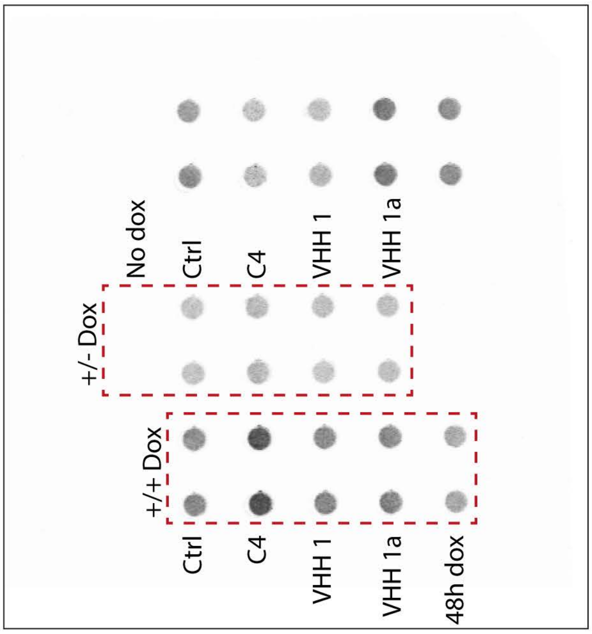
