## Supplementary Tables for "Development of a novel VHH intrabody targeting the N17 region of huntingtin exon 1 protein that prevents inclusion body formation"

Supplementary Table 1. Panning strategy overview.

| **Panning Arm** | **Round 1** | **Round 2** | **Deselection** | **Round 3** | **Competition** | **Deselection** | **Round 4** | **Competition** | **Deselection** |
| --- | --- | --- | --- | --- | --- | --- | --- | --- | --- |
| **HTT**  **alpha** | **N-term N17 peptide biot. [1uM]** | **N-term N17 peptide biot. [500nM]** | C-term HTT peptide biot. | **N-term N17 peptide biot. [200nM]** | N-term N17 peptide [1uM] | C-term HTT peptide biot. | **N-term N17 peptide biot. [50nM]** | N-term N17 peptide [1uM] | C-term HTT peptide biot. |
| **HTT**  **beta** | **PRR peptide biot. [1uM]** | **PRR peptide biot. [500nM]** | C-term HTT peptide biot. | **PRR peptide biot. [200nM]** | PRR peptide [1uM] | C-term HTT peptide biot. | **PRR peptide biot. [50nM]** | PRR peptide [1uM] | C-term HTT peptide biot. |
| **HTT gamma** | **HTT Exon1 Q46 biot. [100nM]** | **HTT Exon1 Q46 biot. [50nM]** | polyQ biot., polyQ biot, C-term HTT peptide biot., MBP | **HTT Exon1 Q46 biot. [10nM]** | HTT Exon1 Q46 [100nM] | polyQ biot., polyP biot., C-term HTT peptide biot., MBP | **HTT Exon1 Q46 biot. [2.5nM]** | HTT Exon1 Q46 [100nM] | polyQ biot., polyP biot, C-term HTT peptide biot., MBP |
| **HTT**  **delta** | **HTT Exon1 Q25 biot.** | **HTT Exon1 Q25 biot.** | HTT Exon1 Q25 biot. | **HTT Exon1 Q25 biot. [10nM]** | HTT Exon1 Q46 [100nM] | HTT Exon1 Q25 biot. | **HTT Exon1 Q25 biot.** | HTT Exon1 Q46 [100nM] | HTT Exon1 Q25 biot. |

Biot = biotinylated. HTT exon 1 Q25 and Q46 were stabilized by fusion of the MBP domain at the N-terminus.

The selection of α-HTT exon 1 specific hVHH binders is divided into four Arms (alpha, beta, gamma, and delta). Binders in Arm alpha are selected toward the N17 N-term peptide with the deselection of the biotinylated C-terminal peptide. Binders in Arm beta are selected toward the PRR peptide with the deselection of the biotinylated C-terminal peptide. Conformational binders in Arm gamma are selected toward the HIS tagged MBP-HTT exon 1 Q46 protein with the deselection of the four biotinylated PolyQ, PolyP, and C-terminal peptides including the HIS tagged MBP protein. Conformational binders in Arm delta are selected toward the HIS-MBP-HTT exon 1 Q46 protein with the deselection of the HIS-MBP-HTT exon 1 Q25 protein. Competition antigen was initiated starting Round 3. The table shows the relevant concentrations used. Starting from Round 2, a step was introduced to deselect irrelevant non-target-specific binders.

Supplementary Table 2. Kinetics KD results using SPR.

| Clone | MBP-HTT exon 1 Q25 | MBP-HTT exon 1 Q46 | N17 peptide | Off target peptide |
| --- | --- | --- | --- | --- |
| VHH 1 | ~207 nM | ~219 nM | ~2.6 µM* | - |
| VHH 2 | ~1010 nM | ~4590 nM | ~16.8 µM* | - |
| VHH 3 | ~112 nM | ~39.5 nM | ~1.6 µM* | - |

* These values are based on fitted values from a single concentration of the peptide and not all three peptide concentrations.

The SPR setup was executed using purified V5-tagged proteins of VHH 1, VHH 2, and VHH 3 captured via the V5 tag onto the chip and assayed for binding to multiple concentrations of the HIS-MBP-HTT exon 1 protein at 33.3, 100, and 300 nM for 200-sec association and 600-sec dissociation time. An off-target VHH clone was included at the end of the run to assess carry-over and cut-off for no-binding (data not shown). The peptide binding was carried out by injection of peptide analytes at 0.12 - 10 μM concentrations at 3-fold dilutions. The association was monitored for 90 sec; dissociation was monitored for 180 sec. Binding kinetics were fitted to a 1:1 binding model after reference and blank subtraction. The off-target C-terminal HTT peptide was taken along as a control.

Supplementary Table 3. Epitope mapping by alanine substitution scan

| N17 peptide variant | VHH1 | VHH2 | VHH3 |
| --- | --- | --- | --- |
| WT | **++** | **++** | **++** |
| N17-M1A | **++** | **-** | **++** |
| N17-T3A | **++** | **-/+** | **++** |
| N17-L4A | **++** | **-** | **++** |
| N17-E5A | **++** | **-** | **++** |
| N17-K6A | **++** | **-/+** | **+** |
| N17-L7A | **++** | **-** | **++** |
| N17-M8A | **-** | **+++** | **++** |
| N17-K9A | **++** | **+** | **+** |
| N17-F11A | **-** | **-/+** | **-** |
| N17-E12A | **-** | **-/+** | **-/+** |
| N17-S13A | **+** | **++** | **++** |
| N17-L14A | **++** | **++** | **++** |
| N17-K15A | **++** | **++** | **+** |
| N17-S16A | **++** | **++** | **++** |
| N17-F17A | **+** | **++** | **++** |

+++ = Max binding (EC50 = 10 – 100 ng/ml); ++ = Good binding (EC50 = 100 – 500 ng/ml); + = relatively good binding (EC50 = 500 – 1000 ng/ml); -/+ = Weak binding (EC50 = 1000 – 10.000 ng/ml); - = No binding.

The table summarizes the EC50 values for binding to each peptide for the Fc-tagged VHH 1, VHH 2, and VHH 3. No binding was observed for all used negative controls (data not shown).

Supplementary Table 4. Epitope mapping by post-translational modification scan

| N17 peptide variant | VHH1 | VHH2 | VHH3 |
| --- | --- | --- | --- |
| WT | **++** | **+** | **++** |
| K6 Acetylation | **++** | **++** | **++** |
| K9 Acetylation | **++** | **++** | **++** |
| K15 Acetylation | **++** | **++** | **++** |
| T3 phosphorylation | **++** | **-** | **++** |
| S13 phosphorylation | **-** | **-/+** | **++** |
| S16 phosphorylation | **++** | **++** | **++** |
| M1 truncation | **++** | **-** | **++** |

+++ = Max binding (EC50 = 10 – 100 ng/ml); ++ = Good binding (EC50 = 100 – 500 ng/ml); + = relatively good binding (EC50 = 500 – 1000 ng/ml); -/+ = Weak binding (EC50 = 1000 – 10.000 ng/ml); - = No binding.

The table provides a summary of the Fc tagged VHH 1, VHH 2, VHH 3 binding to the N17 PTM peptide variants. The data represents the binding of the full-length antibody protein variants to the peptides. No binding was observed for all negative controls (data not shown).
